# Presenilin homologues influence substrate binding and processing by γ-secretase: a molecular simulation study

**DOI:** 10.1101/2023.05.17.541079

**Authors:** Melissa K Eccles, David Groth, Giuseppe Verdile, Mark Agostino

**Affiliations:** Curtin Medical School, Curtin Health Innovation Research Institute, Curtin University, Bentley, Western Australia, Australia; School of Medical and Health Sciences, Edith Cowan University, Joondalup, Western Australia, Australia; Curtin Institute for Computation, Curtin University, Bentley, WA, Australia

**Keywords:** Alzheimer’s disease, beta amyloid, gamma-secretase, metadynamics, presenilin-1, presenilin-2, Notch, amyloid precursor protein

## Abstract

Presenilin homologues in the γ-secretase complex play a pivotal role in substrate binding and processing, impacting β-amyloid (Aβ) peptide generation in Alzheimer’s disease. We conducted a molecular simulation study to determine substrate preferences between presenilin-1 (PS1) and presenilin-2 (PS2) γ-secretase enzymes for amyloid precursor protein (APP) and Notch1 processing. Using homology modelling, we generated PS1- and PS2-γ-secretase models bound to substrates in the Aβ40 and Aβ42 generation pathways and Notch1 S3 and S4 cleavage site substrates. Metadynamics simulations and binding free energy calculations were used to explore conformational ensembles and substrate preferences. PS2-γ-secretase exhibited increased conformational flexibility and preferential binding energy for initiating the Aβ42 pathway compared to PS1-γ-secretase. Additionally, Notch1 exhibits a preference for binding to PS2-γ-secretase over PS1-γ-secretase. This study provides valuable insights into the conformational dynamics of γ-secretase bound to different substrates within a cleavage pathway, improving our understanding of substrate processivity. The findings highlight the importance of considering both PS1- and PS2-γ-secretase in structure-based drug design efforts, with implications for stabilizing or destabilizing specific states during APP processing.

## INTRODUCTION

Alzheimer’s disease (AD) is the most common form of dementia, affecting approximately 35 million people worldwide. With no effective drug interventions that prevent or markedly slow down disease progression, a greater understanding of disease pathogenesis is required.^1^ A key pathological hallmark of AD is the formation of cerebral amyloid plaques, which contain Amyloid-β (Aβ) peptides.^2^ The Aβ peptides are of multiple lengths and are categorized into long (≥42 amino acids) and short (≤40 amino acids) forms, with the long forms being the pathogenic peptides with increased aggregative capacity.^3–5^ The most common Aβ forms generated are Aβ40, accounting for the majority of peptides produced, and Aβ42, accounting for a substantial minority of peptides produced, with the homeostatic Aβ42:Aβ40 ratio being approximately 1:9.^6^ However, these product ratios often shift in AD, such that increased production of longer forms - particularly Aβ42 and Aβ43 - occurs,^3, 7, 8^ leading to increased Aβ aggregation.^4, 5^ Thus, modulation of the production of Aβ peptides represents a potential therapeutic strategy for the treatment of AD.^9, 10^

Aβ peptides are generated via the regulated intramembrane proteolysis (RIP) of **A**myloid **P**recursor **P**rotein (APP), a Type I single-pass transmembrane protein. Initial cleavage by β-APP cleaving enzyme-1 (BACE1) results in ectodomain shedding of APP, allowing the membrane-embedded C-terminal fragment to be cleaved by the integral membrane enzyme γ-secretase, ultimately releasing the **A**PP **I**ntra**C**ellular **D**omain of APP (AICD) and Aβ peptides.^11^ The γ-secretase enzyme (Fig 1A) is a heteromeric multi-subunit integral membrane protein consisting of: Presenilin - the catalytic subunit; Nicastrin (Nct) – involved in substrate gating; Anterior Pharynx Defective 1 (APH1) – a scaffolding and stabilization protein; and Presenilin Enhancer 2 (Pen2) – important for stabilization and enzyme activation.^12–16^ There are two homologues of Presenilin (PS) – Presenilin-1 (PS1) and Presenilin-2 (PS2) – and three isoforms of APH1 – Aph1a^S^, Aph1a^L^, Aph1b – giving rise to six discrete forms of the γ-secretase enzyme.^17^ As the catalytic subunit, the PS homologue has considerable influence over enzymatic activity and genetic mutations in both PS1 and PS2 are causally associated with Autosomal Dominant AD (ADAD).^18–21^

**Figure 1.**
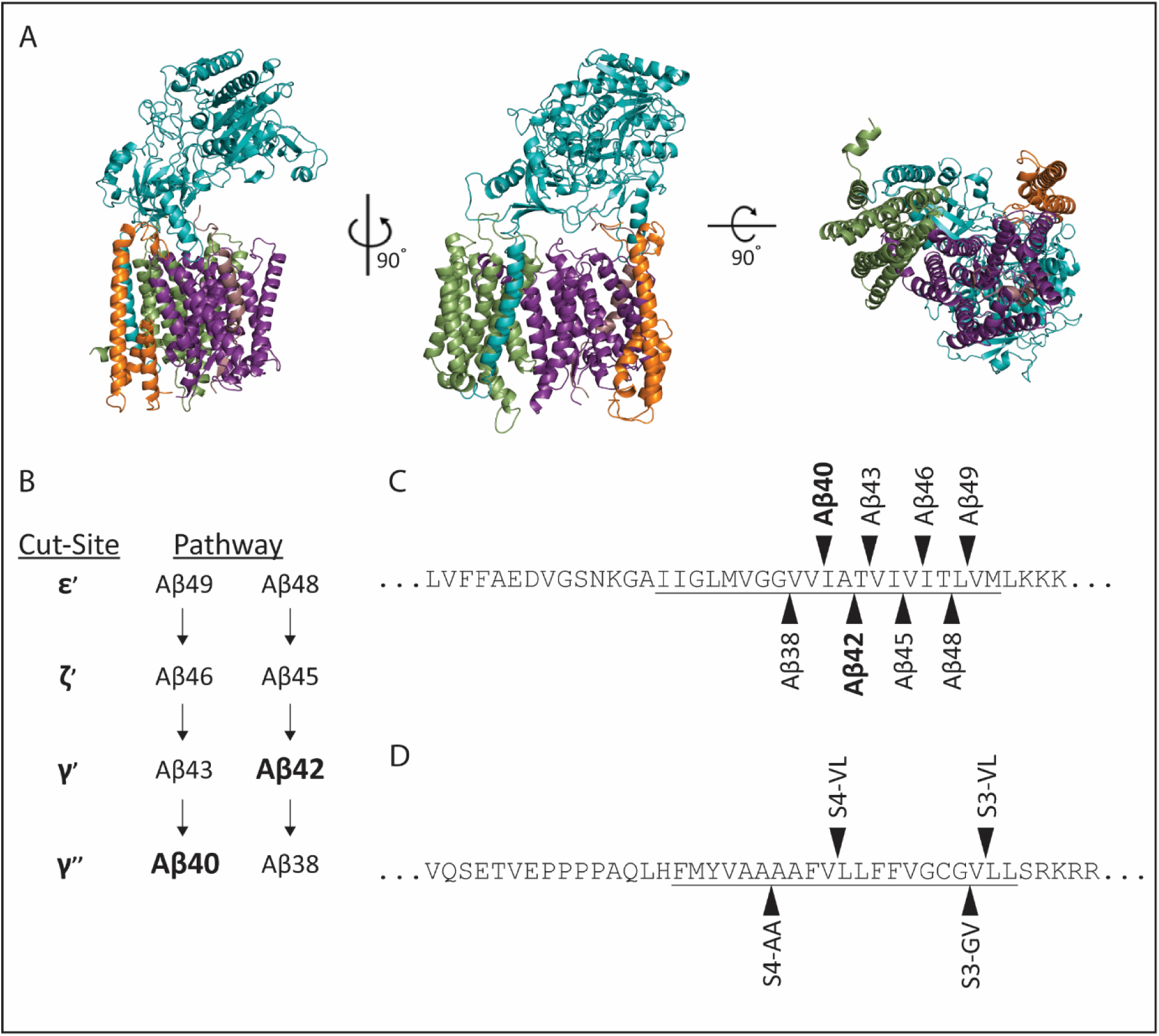
APP and Notch1 processing by γ-secretase. (A) Representative structures of γ-secretase components colored as Aph1a = green, Nicastrin = blue, Pen-2 = orange, Presenilin = purple, APP substrate = light pink. (B) Aβ40 and Aβ42 sequential processing pathways and cut-site terminology.^22^ (C) Position of γ-secretase cleavage sites in APP sequence denoted by arrowhead (π), TMD region of APP-C99 substrate underlined. (D) Position of γ-secretase cleavage sites in Notch1 sequence denoted by arrowhead (π), TMD region of Notch1 substrate underlined.

γ-Secretase generates Aβ peptides via a series of sequential cleavages of the transmembrane domain of APP. These cleavages occur within the lipid bilayer and involve an initial cleavage, releasing the AICD, followed by 2-3 further cleavages spaced at approximately three amino acids between each cleavage (approximately one helical turn). ultimately resulting in the Aβ peptide to be released from the membrane.^22, 23^ The APP processing pathway is well defined, and although there are multiple product lines, two primary pathways have been determined (Fig 1B, C).^23^ These pathways produce Aβ40 and Aβ42 as their major products. The initial substrate, APP-C99, is the same for both product lines, and is cleaved to release the AICD.^22, 23^ However, the cleavage position varies by one residue between the two pathways, with this initial cleavage site being the primary determinant of the product pathway. Cleavage between residues Leu49 and Val50 typically leads to the production of the Aβ40 product, whereas cleavage between Thr48 and Leu49 leads to Aβ42 product generation (Fig 1B, C).^22, 23^

Initial efforts to target γ-secretase involved the development of inhibitors; however, this has been hampered as a therapeutic strategy by side-effects believed to be associated with concurrent inhibition of the processing of other substrates, including Notch-1, which has a diverse array of cell type dependent signaling functions.^24^ Notch-1 is processed similarly to APP, where, prior to γ-secretase processing, ectodomain shedding by ADAM10 occurs,^25, 26^ and the membrane retained product, referred to as the **N**otch **Ex**tracellular **T**runcation (NEXT), is the γ-secretase substrate. NEXT is initially cleaved by γ-secretase at the S3 site to release the **N**otch **I**ntra **C**ellular **D**omain (NICD).^27, 28^ Subsequent cleavage leads to the release of an Aβ-equivalent product, and the final cleavage site is the S4 site (Fig 1D).^28, 29^ Multiple S3 and S4 sites are known, and it is likely that there are similar intermediate processing pathways to those identified for APP processing; however, this has not been definitively determined for Notch. A key advance in improving the functional understanding of γ-secretase has been the determination of its three-dimensional atomic structure via cryo-electron microscopy (cryoEM). The first structure by Bai et al.,^30^ was a 3.4 Å resolution structure (PDB 5A63), followed closely by an additional four structures^31^ ranging from 4.0 – 4.3 Å resolution. These structures included an *apo*-state structure (PDB 5FN5), a DAPT-bound state (a γ-secretase inhibitor; although the structure of the inhibitor could not be resolved) (PDB 5FN2), and two structures containing a volume believed to represent a substrate helix (PDB 5FN3 and 5FN4). Subsequently, higher resolution, substrate-bound cryoEM structures have been published of PS1-γ-secretase bound to APP (2.6 Å) (PDB 6IYC)^32^ and Notch-1 (2.7 Å) (PDB 6IDF)^33^. While these cryoEM structures provide static snapshots, they can be utilized to inform our understanding of the structure and dynamics of γ-secretase. Considerable molecular dynamics work has been completed using these structures, in particular, those prior to the most recent substrate bound structures, investigating the dynamics of substrate docking and entry pathways^34–38^ and the effect of lipids on γ-secretase conformation.^37, 39^ More recently, the effect of presenilin mutations on γ-secretase conformation and substrate binding have been investigated.^40, 41^ Prediction of the binding site of γ-secretase modulators and inhibitors^42, 43^ has been complemented by the most recent atomic structures of PS1-γ-secretase with inhibitory and modulatory small molecules bound.^44^

PS1 is typically considered the more important PS in the context of AD, as the vast majority of pathogenic mutations in the presenilin protein associated with AD occur in PS1 compared with PS2.^45^ However, PS2 expression has been shown to increase over time with *in vitro* neuronal differentiation^46, 47^ and in the human and murine brain with age,^48, 49^ while in cell culture, PS2-γ-secretase has been shown to generate more intracellular Aβ^50, 51^ and increased Aβ42:Aβ40 ratio.^17, 47, 52, 53^ These observations collectively suggest a role for PS2-γ-secretase in disease presentation and progression. Understanding the specific mechanisms and contributions of PS1- and PS2-γ-secretase to substrate binding and processing is therefore important to ensure that effective therapeutics are developed that target PS1 and PS2 complexes effectively.

In this study, we used enhanced sampling approaches, specifically, targeted molecular dynamics and well-tempered metadynamics (WTMetaD), to explore the conformational landscape of PS1- and PS2-γ-secretase in the context of APP and Notch1 processing to examine the similarities and differences and enable improved structural understanding for the future development of Aβ modulating therapies. These approaches were further complemented by binding energy calculations to suggest specific preferences for substrate binding to the different forms of γ-secretase. Traditional experimental methods are limited in their ability to readily provide conformational information about the various states of γ-secretase bound to the multiple Aβ peptides. Similarly, traditional simulation approaches may overly sample specific protein conformations and lead to an incomplete understanding of the potential states that may be adopted by a given protein or protein complex. Homology modelling and metadynamics provide a tool kit that can allow the investigation of multiple γ-secretase and substrate combinations to understand the conformational ensembles of PS1-vs PS2-γ-secretase processing of APP and Notch1.

## RESULTS

### Derivation of path between APP-bound and Notch1-bound PS1-γ-secretase

Targeted molecular dynamics and analysis of the derived trajectories was initially performed to identify a potential path linking the APP-bound and Notch1-bound states of PS1-γ-secretase. The path ultimately identified was 20 frames in length. Key regions of dynamic movement in the frames comprising the path, representing key regions of structural variation between the two complex structures, are the region between the second and third transmembrane helices of presenilin, the region between the sixth and seventh transmembrane helices of presenilin, and the region between the third and fourth transmembrane helices of Aph1a (Fig S1A). All of these regions are located on the lumenal side of the complex. The corresponding PS2-γ-secretase path was derived via homology modelling to the component frames of the PS1-γ-secretase path, on the assumption that PS2-γ-secretase is likely to display similar dynamics to PS1-γ-secretase.

### WTMetaD of PS1- and PS2-γ-secretase bound to APP substrates

Using the cryoEM structure of PS1-γ-secretase bound to APP-CTF in position for the initial cleavage of the Aβ40 pathway (PDB:6IYC)^32^, homology models of PS1 and PS2-γ-secretase bound to APP-CTF (and intermediate fragments) in position for cleaving Aβ49, Aβ48, Aβ46, Aβ45, Aβ43, Aβ42, Aβ40, and Aβ38 were prepared. Using the path derived from targeted MD simulation, WTMetaD in the position along (denoted as *s*, where *s* = 1 represents γ-secretase bound to APP as per PDB 6IYC, herein referred to as the 6IYC-like conformation of γ-secretase, and *s* = 20 represents γ-secretase bound to Notch1 as per PDB 6IDF, herein referred to as the 6IDF-like conformation of γ-secretase) and deviation from (denoted as *z*, with *z = 0* being along the path, and *z* > 0 being a deviation from the path) the path was performed. WTMetaD of both PS1- and PS2-γ-secretase bound to the initial APP-CTF substrate in the cleavage position for initiating the Aβ40 pathway (i.e. to generate the intermediate Aβ49 product) revealed only one major energetic minimum for both complexes, with similar 6IYC-like conformations (Fig 2A: PS1:CTF(Aβ49) *s* ≈ 4, *z ≈* 0.04; Fig 2B: PS2:CTF(Aβ49) *s* ≈ 3, *z* ≈ 0.035).

**Figure 2.**
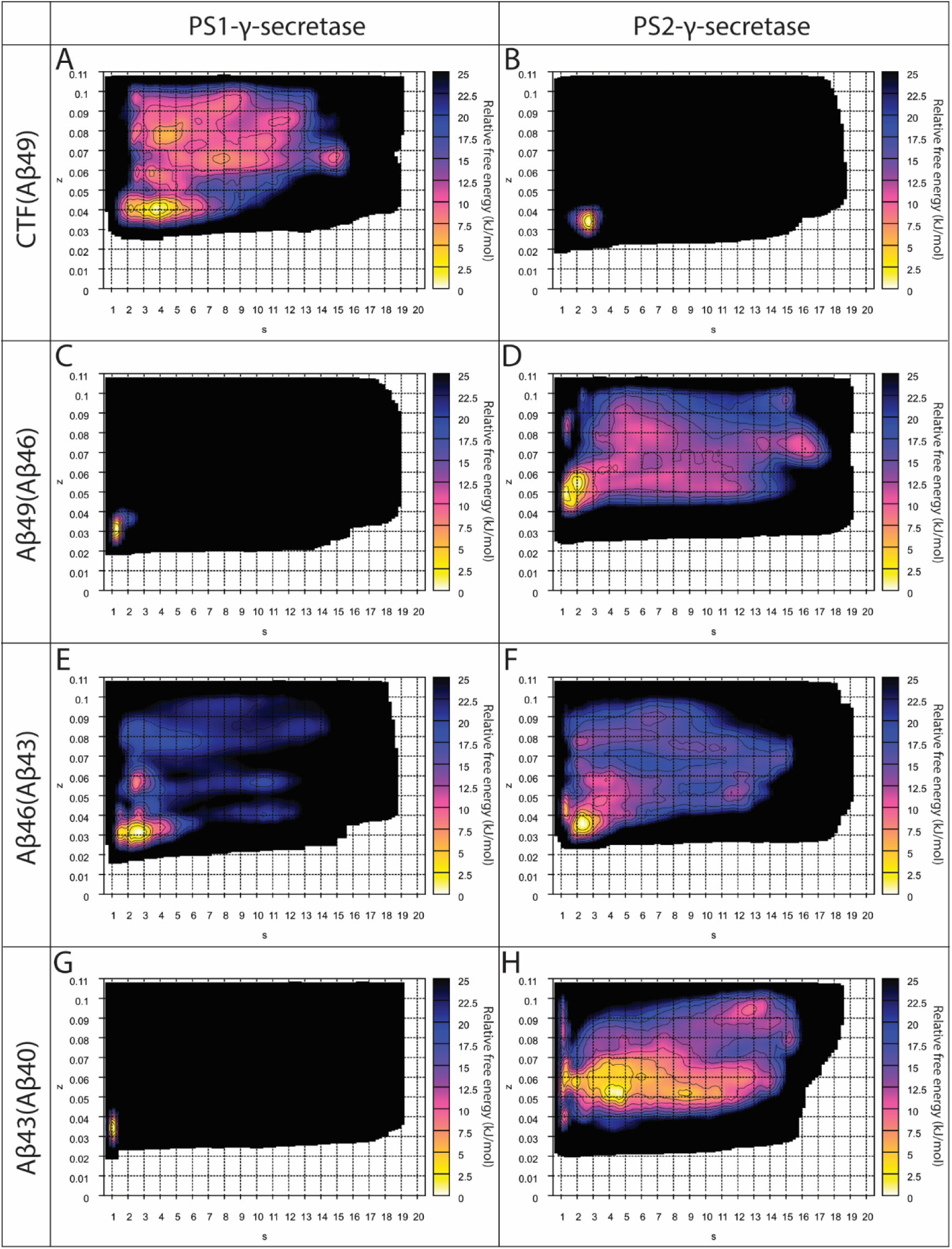
Well-tempered metadynamics simulations of PS1- and PS2-γ-secretase in complex with Aβ40 pathway substrates. Relative free energy surfaces for (A, C, E, G) PS1-γ-secretase and (B, D, F, H) PS2-γ-secretase in complex with substrates [denoted as substrate(cleavage position)] (A, B) CTF(Aβ49), (C, D) Aβ49(Aβ46), (E, F) Aβ46(Aβ43), (G, H) Aβ43(Aβ40).

Energetic minima derived from WTMetaD of PS1- and PS2-γ-secretases bound to APP-CTF in the cleavage position to initiate the Aβ42 pathway (i.e., to generate the intermediate Aβ48 product) revealed two energetic minima for the PS1:CTF(Aβ48) complex, both of which are 6IYC-like (Fig 3A: PS1:CTF(Aβ48) *s* ≈ 3, *z ≈* 0.035 and *s* ≈ 1.5, *z ≈* 0.08). The lowest energy minimum of the PS1:CTF(Aβ48) complex (*s* ≈ 3, *z ≈* 0.035) is comparable to the PS1:CTF(Aβ49) complex minimum in contour breadth, suggesting a similar conformational flexibility in PS1-γ-secretase when binding either substrate. However, the PS2 complex has only one 6IYC-like energetic minimum (Fig 3E: PS2:CTF(Aβ48) *s* ≈ 1.5, *z ≈* 0.04), and while similarly positioned with respect to the path compared to the PS2:CTF(Aβ49) complex, this minimum has broader contours, suggesting that this complex is likely more flexible than the equivalent complex with PS1.

**Figure 3.**
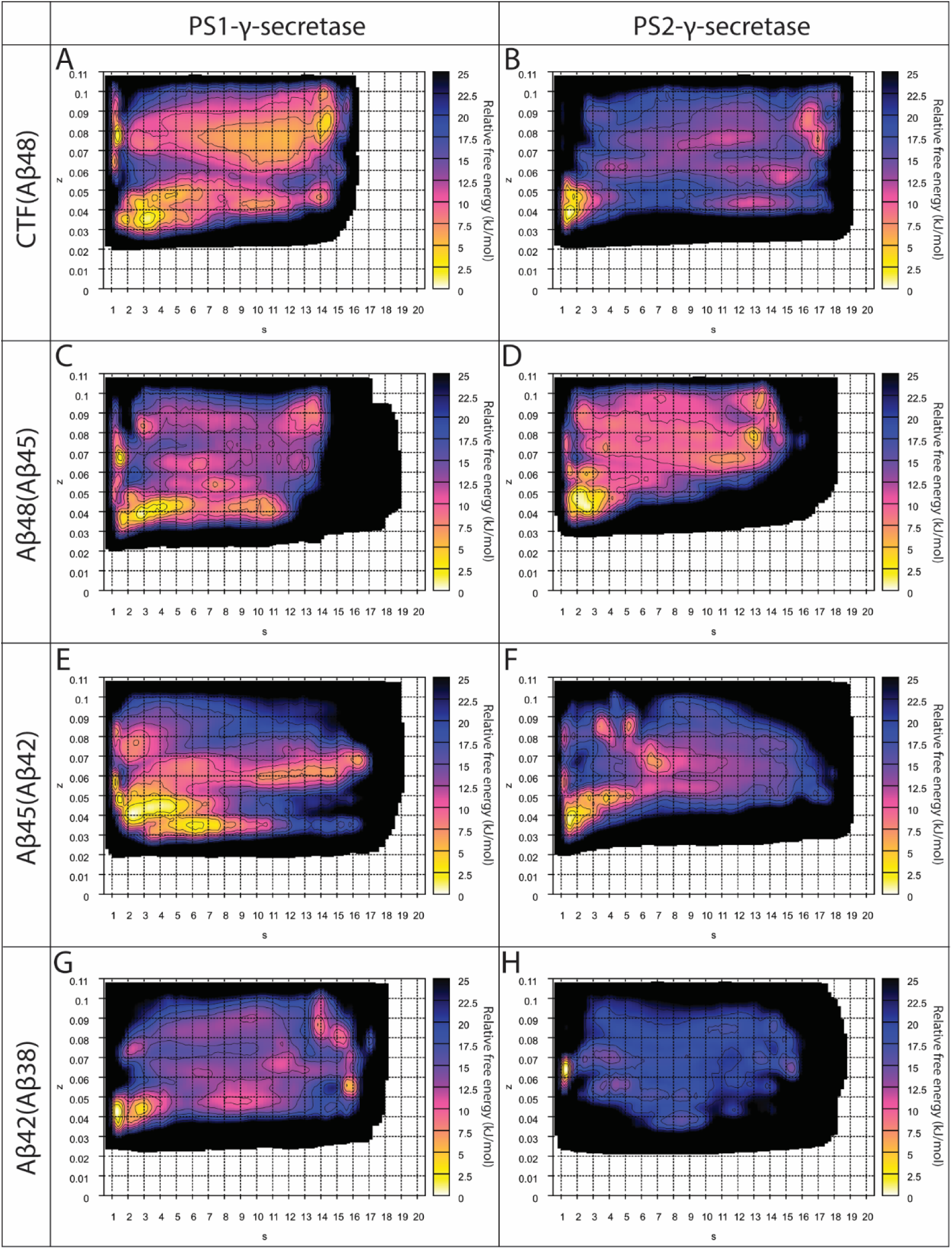
Well-tempered metadynamics simulations of PS1- and PS2-γ-secretase in complex with Aβ42 pathway substrates. Relative free energy surfaces for (A, C, E, G) PS1-γ-secretase and (B, D, F, H) PS2-γ-secretase in complex with substrates [denoted as substrate(cleavage position)] (A, B) CTF(Aβ48), (C, D) Aβ48(Aβ45), (E, F) Aβ45(Aβ42), (G, H) Aβ42(Aβ38).

WTMetaD of PS1- and PS2-γ-secretase in complex with all intermediate substrates of the Aβ40 and Aβ42 processing pathways was then performed (Fig 2 & 3). All subsequent γ-secretase– substrate complexes, after the initial cleavage to release AICD, have one energetic minimum, with the exception of the PS1:Aβ48(Aβ45) complex, which has two energetic minima (Fig 3B). All minima are 6IYC-like (*s* < 5 in all cases). The contour breadth in the PS1-γ-secretase complexes with Aβ40 pathway substrates is considerably tighter in the shorter substrates further along the pathway, in comparison to the initial CTF(Aβ49) substrate (Fig 2B-D), compared to the broader contours in the equivalent complexes with PS2-γ-secretase (Fig 2F-H). In contrast, PS2-γ-secretase displays a restricted conformational ensemble when binding to the initial CTF(Aβ48) substrate in the Aβ42 pathway (Fig 3E), while subsequent enzyme-substrate complexes in the pathway (with the exception of the PS2:Aβ42(Aβ38) complex) display relatively broader free energy surface contouring, suggesting increased conformational flexibility (Fig 3F-H).

### Binding free energies of γ-secretase - APP bound complexes

Molecular mechanics-generalized Born/surface area calculations (MM-GB/SA) were performed for all enzyme-substrate complexes to estimate the binding energies (ΔG_bind_) for γ-secretase with each of the substrates (Table 1). Structures corresponding to low energy regions of the free energy surfaces derived from the WTMetaD simulations were used for the MM-GB/SA calculations. The PS1:CTF(Aβ49) complex, in the position to initiate the Aβ40 pathway, has a more favorable binding energy compared to the PS2:CTF(Aβ49) complex. However, the opposite preference is observed for CTF(Aβ48), where PS2:CTF(Aβ48) has a marginally lower binding energy than PS1:CTF(Aβ48). While there are two minima evident from the metadynamics simulation for the PS1:CTF(Aβ48) complex, MM-GB/SA calculations at both minima afford a higher binding energy compared to the PS2:CTF(Aβ48) complex. Thus, CTF(Aβ48) is predicted to have a preference for binding PS2-γ-secretase over PS1-γ-secretase. Considering the binding energy results for all four complexes bound to the APP-CTF, there is a preference for both PS1- and PS2-γ-secretase to bind the APP-CTF in the Aβ48 position over the Aβ49 position.

**Table 1.** MM-GB/SA free binding energy of γ-secretase – APP bound complexes.

| Aβ40 Pathway | | | | Aβ42 Pathway | | | |
| --- | --- | --- | --- | --- | --- | --- | --- |
| Substrate | Product | PS1<sup>§</sup> | PS2<sup>§</sup> | Substrate | Product | PS1<sup>§</sup> | PS2<sup>§</sup> |
| APP-CTF | Aβ49 | -189.7 ± 1.5 | -178.4 ± 2.7 | APP-CTF | Aβ48 | -205.6 ± 1.8* | -208.9 ± 2.6 |
| Aβ49 | Aβ46 | -142.2 ± 3.1 | -158.3 ± 1.1 | Aβ48 | Aβ45 | -181.1 ± 0.8<sup>#</sup> | -164.7 ± 1.8 |
| Aβ46 | Aβ43 | -165.0 ± 3.4 | -148.6 ± 1.3 | Aβ45 | Aβ42 | -145.6 ± 0.5 | -122.4 ± 1.5 |
| Aβ43 | Aβ40 | -123.2 ± 1.9 | -124.4 ± 1.3 | Aβ42 | Aβ38 | -153.1 ± 1.1 | -132.3 ± 1.0 |

\*The PS1-γ-secretase complex bound to APP-CTF in position to generate the Aβ48 product has a second minima located at $s = 0$ to $s = 2$ and $z = 0.07$ to $z = 0.09$ , this complex conformation has a MMGBSA free binding energy of -194.7 ± 3.0 kCal/mol
<sup>#</sup>The PS1-γ-secretase complex bound to Aβ48 to generate the Aβ45 product has a second minima located at $s = 0$ to $s = 3$ and $z = 0.06$ to $z = 0.08$ , this complex conformation has a MMGBSA free binding energy of -181.1 ± 3.8 kcal/mol
<sup>§</sup>Values shown are in kcal/mol +/- standard error

The binding energies for the subsequent complexes in the Aβ40 pathway suggest that while PS2-γ-secretase has a preference for binding to the Aβ49(Aβ46) substrate over PS1-γ-secretase, the PS2-γ-secretase enzyme binds Aβ46(Aβ43) significantly less favorably than the PS1-γ-secretase enzyme. Both PS1- and PS2-γ-secretase enzymes have approximately equal preference for binding the Aβ43(Aβ40) substrate. In the Aβ42 pathway, however, the subsequent PS1 complexes - PS1:Aβ48(Aβ45), PS1:Aβ45(Aβ42) and PS1:Aβ42(Aβ38) – have consistently more favorable binding energy than the equivalent PS2 complexes.

### Per residue decomposition of binding free energies for γ-secretase-APP bound complexes

Per-residue decompositions of the MM-GB/SA binding energies were performed to determine the individual contributions of residues from the enzyme and the substrate to the overall binding energy. The relative change in binding energy per-residue with respect to PS1-versus PS2-γ-secretase was also calculated (ΔΔG_PS Pref_, calculation described in Methods) to determine the precise contributors to selectivity (Figures 4 and 5).

**Figure 4.**
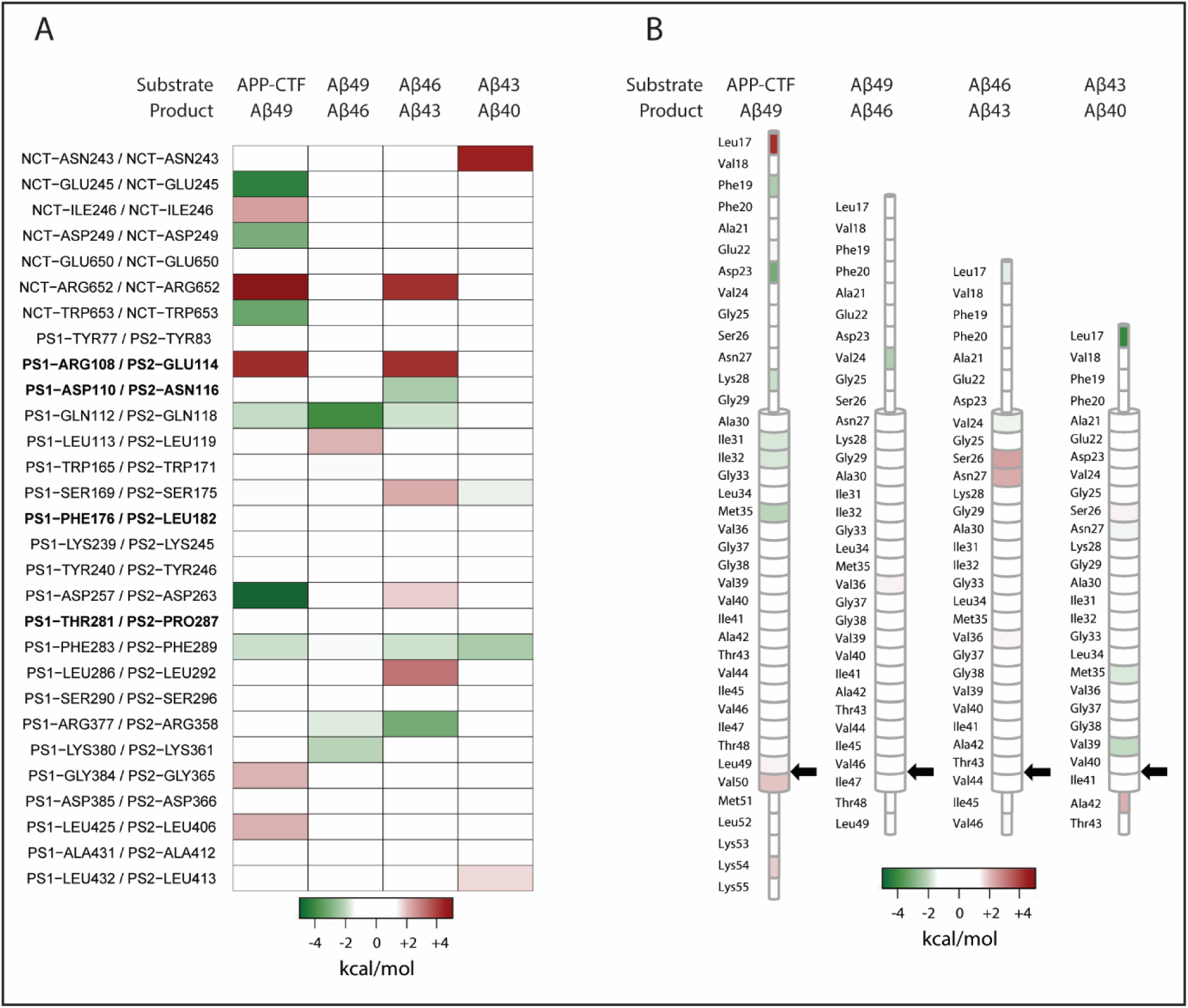
Aβ40 pathway complexes per residue heatmaps of ΔΔGPS Pref. (A) Enzyme ΔΔGPS Pref values. Only residues where the ΔΔGPS Pref magnitude is greater than 1.5 kcal/mol in any complex are shown. (B) Substrate ΔΔGPS Pref values. Cleavage position denoted by arrow. Positive (red) ΔΔGPS Pref values indicate preference for PS1, negative (green) ΔΔGPS Pref values indicate preference for PS2.

**Figure 5.**
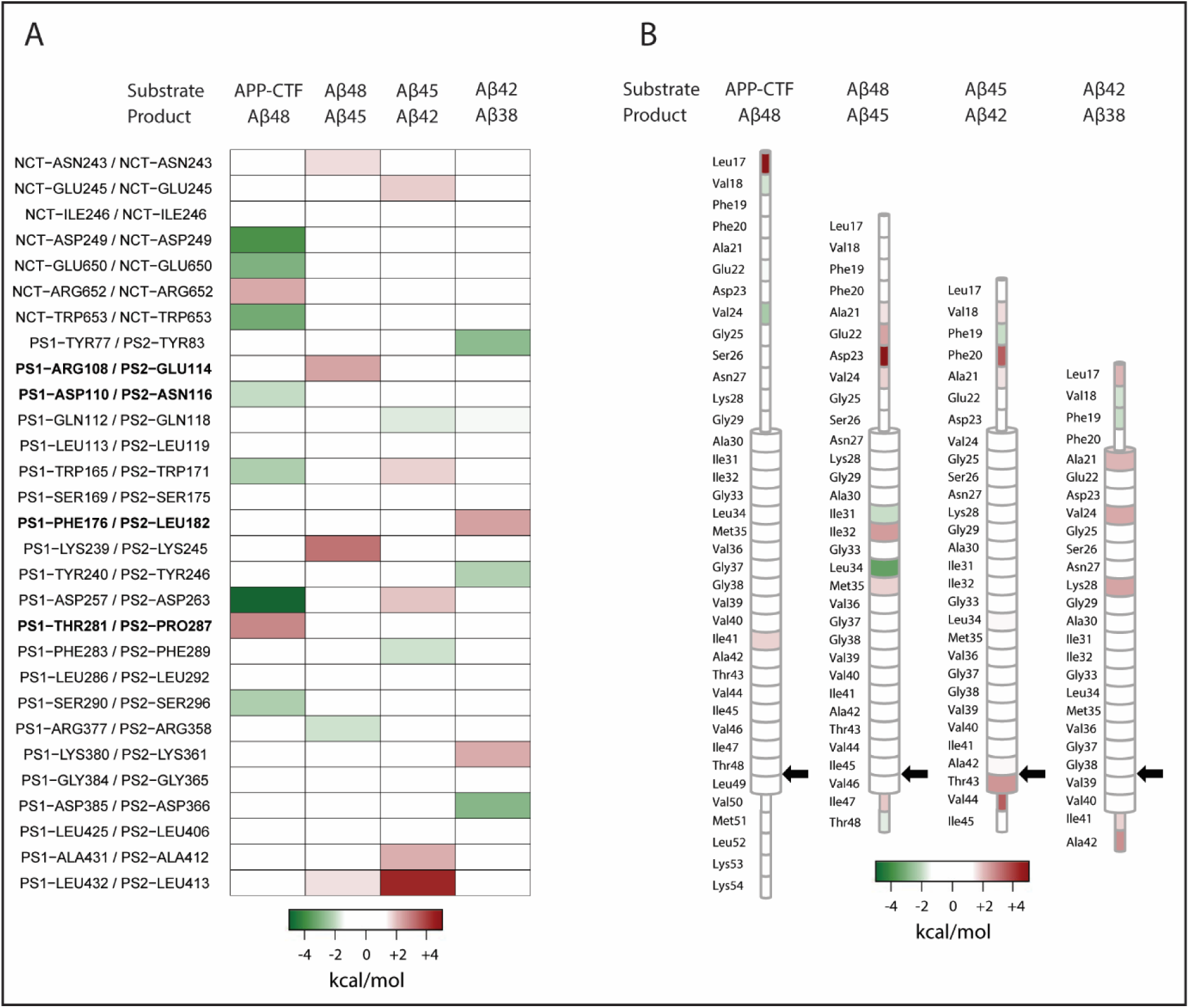
Aβ42 pathway complexes per residue heatmaps of ΔΔGPS Pref. (A) Enzyme ΔΔG PS Pref values. Only residues where the ΔΔGPS Pref magnitude is greater than 1.5 kcal/mol in any complex are shown. (B) Substrate ΔΔGPS Pref values. Cleavage position denoted by arrow. Positive (red) ΔΔGPS Pref values indicate preference for PS1, negative (green) ΔΔGPS Pref values indicate preference for PS2.

The APP-CTF substrate bound in position to produce the Aβ49 product, thus initiating the Aβ40 pathway, shows preference for binding to PS1-γ-secretase over PS2-γ-secretase. This preference is driven by a series of contributions clustered at the N-terminus of the substrate, specifically, nicastrin residues Ile246 (ΔΔG_PS Pref_ = +2.49 kcal/mol) and Arg652 (ΔΔG_PS Pref_ = +6.07 kcal/mol), which form hydrogen bonds (mediated by the Ile246 backbone amide) and a salt bridge respectively with Glu22 (ΔG_bind PS1_ = +0.87 kcal/mol) in the substrate. Substrate residue Leu17 (ΔΔG_PS Pref_ = +4.26 kcal/mol), which forms a hydrogen bond between its backbone carbonyl with the backbone amide of Trp653 (ΔG_bind PS1_ = −8.64 kcal/mol) in nicastrin, and the PS1 residue Arg108 (ΔΔG_PS Pref_ = +4.29 kcal/mol), which is likely forming cation-π interactions with the substrate residues Phe19 (ΔG_bind PS1_ = −4.60 kcal/mol) and Phe20 (ΔG_bind PS1_ = −2.17 kcal/mol), also contribute to this preference. Arg108 also forms a salt bridge with Nct Glu245. Arg108 is not conserved in PS2 and is replaced by a glutamate at the analogous position (PS2 Glu114), with this residue unable to make the same interactions (Fig 4 & S2A). Further contributions more favorable in the PS1-γ-secretase complex occur at the C-terminal region of the substrate, specifically involving Val50 (ΔΔG_PS Pref_ = +1.94 kcal/mol), which forms multiple hydrophobic interactions with Val272 (ΔG_bind PS1_ = −1.22 kcal/mol) and Ile287 (ΔΔG_PS Pref_ = −1.47 kcal/mol) in PS1, and Lys54 (ΔΔG_PS Pref_ = +1.86 kcal/mol), which forms hydrogen bonds and a salt bridge between PS1 residues Thr291 (ΔΔG_PS Pref_ = +0.56 kcal/mol) and Glu376 (ΔΔG_PS Pref_ = +0.99 kcal/mol), respectively (Fig 4 & S2B). Additionally, PS1 residue Gly384 (ΔΔG_PS Pref_ = +2.16 kcal/mol) forms hydrogen bonds with the backbone of Val46 (ΔΔG_PS Pref_ = +0.13 kcal/mol) and PS1 Leu425 (ΔΔG_PS Pref_ = +2.18 kcal/mol) hydrophobically interacts with substrate residue Met51 (ΔΔG_PS Pref_ = +0.77 kcal/mol). These interactions likely contribute to the stabilization of the β-sheet structure between PS1 and the substrate that is a key feature of the γ-secretase bound substrate,^32, 33^ and do not readily occur in the PS2 complex.

The subsequent complex in the Aβ40 pathway, the Aβ49 substrate positioned to produce the Aβ46 product, demonstrates a preference for binding to PS2-γ-secretase over PS1-γ-secretase. This preference is a result of two key regions of interactions. PS2 Gln118 (ΔΔG_PS Pref_ = −3.97 kcal/mol) forms a hydrogen bond with substrate residue Val24 (ΔΔG_PS Pref_ = −2.19 kcal/mol) in the luminal juxtamembrane region, which does not occur in the PS1-complex (Fig 4 & S2C). The PS2 residues Arg358 (ΔΔG_PS Pref_ = −1.64 kcal/mol) and Lys361 (ΔΔG_PS Pref_ = −2.08 kcal/mol) coordinate a network of hydrogen bonds with the substrate C-terminal residues Thr48 (ΔΔG_PS Pref_ = −0.26 kcal/mol) and Leu49 (ΔΔG_PS Pref_ = −0.76 kcal/mol), likely stabilizing the C-terminus of the substrate; the majority of these interactions do not occur in the PS1-complex (Fig 4 & S2D).

The Aβ46 substrate, processing of which leads to the generation of the Aβ43 product in the Aβ40 pathway, shows a preference for binding to PS1-γ-secretase over PS2-γ-secretase. Key contributors to the binding preference are the nicastrin residue Arg652 (ΔΔG_PS Pref_ = +4.27 kcal/mol), and the PS1 residue Arg108 (ΔΔG_PS Pref_ = +4.24 kcal/mol), which forms electrostatic interactions with Glu22 (ΔG_bind PS1_ = −0.85 kcal/mol) in the substrate. As previously noted, the PS1 Arg108 residue is not conserved in PS2, with the analogous residue in PS2 being Glu114; consequently, these interactions cannot occur in the PS2-complex. Substrate residues at the N-terminal juxtamembrane region, Ser26 (ΔΔG_PS Pref_ = +2.45 kcal/mol) and Asn27 (ΔΔG_PS Pref_ = +2.28 kcal/mol), further contribute to the substrate preference for PS1 by forming a greater network of hydrogen bonds with residues in the loop between TM-1 and TM-2 (Ile114: ΔΔG_PS Pref_ = +0.81 kcal/mol; Thr116: ΔΔG_PS Pref_ = +0.58 kcal/mol), which does not occur in the PS2-complex (Fig 4 & S2E). Additionally, a series of interactions between PS1 and substrate residues in the inner-leaflet transmembrane domain (TMD) region and the C-terminal juxta-membrane region form, that do not occur in the PS2-complex (Fig 4 & S2F). PS1 residue Ser169 (ΔΔG_PS Pref_ = +2.28 kcal/mol) forms hydrogen bonds between its backbone amine group and the backbone carbonyl group of Met35 (ΔG_bind PS1_ = −1.51 kcal/mol), as well as utilizing its sidechain hydroxyl group to form a hydrogen bond with the amide of Gly38 (ΔΔG_PS Pref_ = +0.54 kcal/mol). Leu286 (ΔΔG_PS Pref_ = +3.17 kcal/mol) forms a network of hydrophobic interactions with Ile41 (ΔΔG_PS Pref_ = +0.80 kcal/mol) and Ala42 (ΔΔG_PS Pref_ = +0.71 kcal/mol); notably, these appear to stabilize the β-sheet structure between the substrate C-terminus and PS1 previously noted.^32, 33^ These interactions occur in tandem with the Trp165 (ΔG_bind PS1_ = - 4.28 kcal/mol) sidechain forming a CH-π bond with Gly38 (ΔΔG_PS Pref_ = 0.54 kcal/mol) in the substrate. The analogous tryptophan in PS2, Trp171 (ΔG_bind PS2_ = −4.29 kcal/mol), is positioned such that it interacts with Val39 (ΔG_bind PS2_ = −4.61 kcal/mol), forming a CH-π bond. The residue adjacent to Trp165/Trp171 is not conserved between PS1 and PS2, being an alanine in PS1 (Ala164) and a glycine in PS2 (Gly170); this likely influences the presentation of the tryptophan sidechain, as the backbone in this region is expected to exhibit greater flexibility in PS2-vs PS1-γ-secretase.

APP-CTF bound in the position to generate the Aβ48 product, representing the initial substrate in the Aβ42 pathway, has a slight preference for binding to PS2-γ-secretase over PS1-γ-secretase. The primary residue contributing to this preference is the PS2 residue Asp263 (ΔΔG_PS Pref_ = −9.28 kcal/mol), which is the catalytic aspartate in the N-terminal fragment of PS2 (Fig 5). Asp366, protonated in our models, is presented in PS2 in a manner that allows for multiple hydrogen bonds between its side chain with substrate residue Thr48 and the backbone amine of Leu49 (Fig 5 & S3A), unlike the equivalent residue in PS1, where these interactions are absent. Additionally, the N-terminal residues of APP-CTF (Val18 (ΔΔG_PS Pref_ = −1.75 kcal/mol) and Phe20 (ΔΔG_PS Pref_ = −0.24 kcal/mol) interact with a cluster of nicastrin residues (Asp249 (ΔΔG_PS Pref_ = −3.80 kcal/mol), Glu650 (ΔΔG_PS Pref_ = −2.98 kcal/mol), and Trp653 (ΔΔG_PS Pref_ = −3.18 kcal/mol)) and residues in the TM1 to TM2 loop region of PS2 (Asn116 (ΔΔG_PS Pref_ = −1.94 kcal/mol)), forming hydrogen bonds and hydrophobic interactions that are not evident in the PS1-γ-secretase complex (Fig 5 & S3B). Notably, Asn116 in PS2 is not conserved in PS1, the analogous residue being Asp110; the negatively charged aspartate residue in PS1 forms a salt-bridge with the positively charged Arg108 in PS1, preventing interactions with the substrate.

The subsequent substrates in the Aβ42 pathway all demonstrate preference for binding to PS1-γ-secretase (Table 1). The primary contributors to this preference in the γ-secretase complexes bound to the Aβ48 substrate in position to generate the Aβ45 product are the nicastrin residue Asn243 (ΔΔG_PS Pref_ = +1.68 kcal/mol), and the PS1 residues Arg108 (ΔΔG_PS Pref_ = +2.41 kcal/mol) and Lys239 (ΔΔG_PS Pref_ = +3.15 kcal/mol). These enzyme residues are positioned around a cluster of residues in the N-terminal juxtamembrane region of the substrate, including Ala21 (ΔΔG_PS Pref_ = +1.62 kcal/mol), Glu22 (ΔΔG_PS Pref_ = +2.39 kcal/mol), Asp23 (ΔΔG_PS Pref_ = +5.13 kcal/mol), and Val24 (ΔΔG_PS Pref_ = +1.76 kcal/mol), forming a salt bridge and hydrogen bonds (Fig 5 & S3C). This cluster of interactions is facilitated by PS1 Arg108, which interacts with the main chain carbonyl groups of the substrate residues Ala21 and Glu22, forming hydrogen bonds and stabilizing the substrate N-terminus. These interactions are not formed in the PS2 complex, as Arg108 is replaced with Glu114 in PS2 (Fig 5 & S3C). Additionally, a network of hydrophobic interactions forms between substrate residue Ile32 (ΔΔG_PS Pref_ = +2.53 kcal/mol), with PS1 Ile114 and Tyr240, and Met35 (ΔΔG_PS Pref_ = +1.81 kcal/mol), with PS1 Leu172, Phe177 and Val236 (Fig 5 & S3D). These interactions do not form in the PS2-complex; notably, PS1 residue Leu172 is not conserved (analogous PS2 residue is Met178), nor is the residue immediately adjacent to Phe177 (PS1 Phe176 analogous PS2 residue is Leu182), likely influencing the observed interactions between the PS1- and PS2-complex. The substrate helix itself in the PS2-complex is observably disrupted at its di-glycine motif, which has been identified as a point of flexibility in APP,^54^ and also likely contributes to the reduced interactions with PS2 residues in the substrate binding pocket. Lastly, interactions involving Ile47 (ΔΔG_PS Pref_ = +1.91 kcal/mol), at the C-terminal end of the substrate, which interacts via hydrogen bonds to the backbone and hydrophobic interactions to the sidechain of Leu432 (ΔΔG_PS Pref_ = +1.63 kcal/mol), are not replicated in the PS2-complex.

The Aβ45 substrate, which is cleaved to the Aβ42 product, similarly shows a preference for binding to PS1-γ-secretase over PS2-γ-secretase. Key residues contributing to this preference include the nicastrin residue Glu245 (ΔΔG_PS Pref_ = +1.84 kcal/mol), which is positioned proximally to a cluster of residues in the N-terminal juxta-membrane region that contribute to the preference for PS1-γ-secretase, specifically, Val18 (ΔΔG_PS Pref_ = +1.72 kcal/mol), Phe20 (ΔΔG_PS Pref_ = +3.54 kcal/mol), Ala21 (ΔΔG_PS Pref_ = +1.56 kcal/mol) (Fig 5 & S3E). The N-terminal region of the substrate appears to be stabilized by cation-π interactions between PS1 Arg108 and substrate residue Phe20; equivalent interactions do not occur in the PS2-complex as the analogous residue to Arg108 in PS2 is the negatively charged Glu114. The primary PS1 residues that contribute to this preference are Ala431 (ΔΔG_PS Pref_ = +2.23 kcal/mol), which forms hydrophobic interactions with Ile45, and the adjacent Leu432 (ΔΔG_PS Pref_ = +4.36 kcal/mol), which forms hydrogen bonds between the backbone amide group and the carbonyl of Val44 (ΔΔG_PS Pref_ = +3.45 kcal/mol) in the substrate. Additionally, the substrate residue Thr43 (ΔΔG_PS Pref_ = +2.55 kcal/mol) contributes to the PS1-complex preference via hydrophobic interactions between its side chain carbon and the sidechains of PS1 Val261 and Val272. The β-strand of the substrate contributing to the hybrid β-sheet with presenilin is disrupted in the PS2-complex, which appears to preclude the majority of these interactions from occurring (Fig 5 & S3F).

The final complexes modelled in the Aβ42 pathway feature Aβ42 itself as the substrate, positioned for the generation of Aβ38 as the final product in the pathway. From the binding free energies determined by MM-GB/SA, this substrate exhibits a preference for PS1-γ-secretase over PS2-γ-secretase. The preference is driven by the PS1 residue Phe176 (ΔΔG_PS Pref_ = +2.44 kcal/mol), which can form NH-π interactions^55^ with the substrate residue Asn27. An equivalent interaction is unable to form in the PS2 complex, as Phe176 is replaced by Leu182 (Fig S3G). Substrate residues Val24 (ΔΔG_PS Pref_ = +2.33 kcal/mol) and Lys28 (ΔΔG_PS Pref_ = +2.37 kcal/mol) (specifically, carbons along the sidechain) interact with Tyr240 (ΔG_bind PS1_ = - 1.51 kcal/mol) and Ile114 (ΔΔG_PS Pref_ = +0.98 kcal/mol) in PS1, respectively, via hydrophobic interactions (Fig 5 & S3G). Additionally, the substrate residues Ile41 (ΔΔG_PS Pref_ = +1.76 kcal/mol) and Ala42 (ΔΔG_PS Pref_ = +2.73 kcal/mol) form a spatial cluster of residues at the C-terminus of the substrate that contribute to the PS1 complex preference. This cluster of interactions is predominated by salt-bridge interactions between PS1 Lys380 (ΔΔG_PS Pref_ = +2.26 kcal/mol) and Lys429 (ΔΔG_PS Pref_ = +1.45 kcal/mol) with the carboxyl group on the C-terminus of the substrate, interactions that do not occur in the modelling of the PS2 complex (Fig 5 & S3H).

### WTMetaD and binding free energy calculations for PS1- and PS2-γ-secretase bound to Notch1 substrates

Given the functional implications of Notch1 inhibition, it is imperative that Notch1 processing by γ-secretase is considered in future therapeutic targeting of the enzyme. Consequently, we examined Notch1 bound to PS1- and PS2-γ-secretase enzymes positioned for processing at the two primary S3 sites - Val1754/Leu1755 and Gly1753/Val1754 - and the two primary S4 sites - Val1745/Leu1746 and Ala1741/Ala1742. The cryoEM structure of PS1-γ-secretase bound to Notch1 in the Val1754/Leu1755 (PDB: 6IDF) ^33^ was used to generate homology models of PS1- and PS2-γ-secretase enzymes bound to substrates in the S3 and S4 positions.

WTMetaD in the position along and deviation from the path was performed, using the path derived from targeted MD. WTMetaD of all four substrate/cleavage positions in complex with the PS1- or PS2-γ-secretase enzyme revealed only one energetic minimum for each complex, except for PS1-γ-secretase bound to the S3 substrate in position to cleave at the Ala1741/Ala1742 S4 site, where two minima are evident (Fig 6). The energetic minima derived from the PS2-γ-secretase bound to NEXT in the two S3 cleavage positions are both in a more 6IDF-like position (PS2:NEXT(VL) *s ≈ 14, z ≈ 0.04*; PS2:NEXT(GV) *s ≈ 15, z ≈ 0.04*) compared with the PS1-complexes (PS1:NEXT(VL) *s ≈ 12, z ≈ 0.04*; PS2:NEXT(GV) *s ≈ 11, z ≈ 0.04*). The resultant S3 product, after the cleavage and release of the NICD, was then examined. The energetic minima for the substrate positioned for either of the S4 cleavages are similar (PS1:S3(VL) *s ≈ 14, z ≈ 0.04*; PS2:S3(VL) *s ≈ 13, z ≈ 0.04,* PS1:S3(AA) *s ≈ 12, z ≈ 0.05* and *s ≈ 14, z ≈ 0.05*; PS2:S3(AA) *s ≈ 14, z ≈ 0.04*).

**Figure 6.**
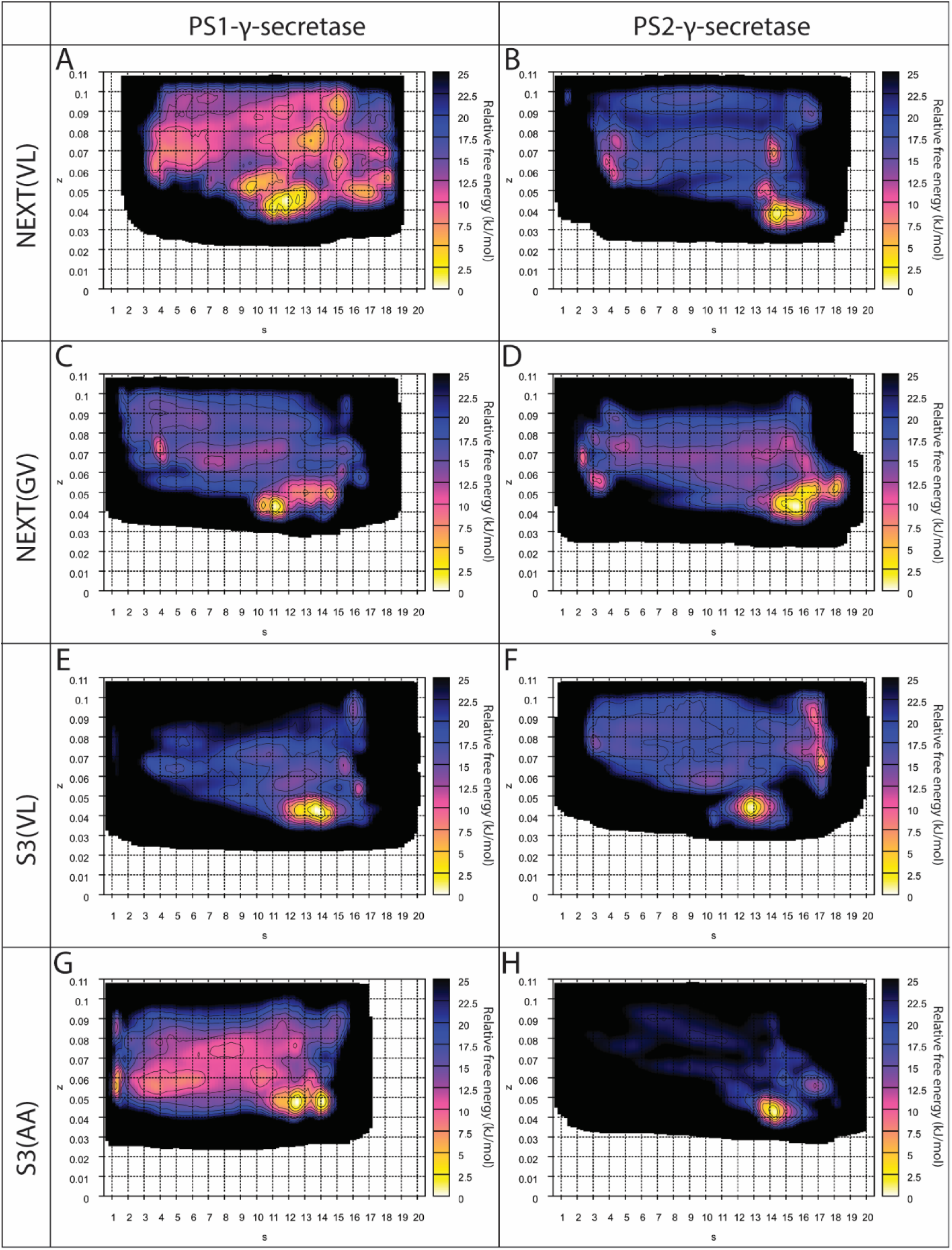
Well-tempered metadynamics simulations of PS1- and PS2-γ-secretase in complex with Notch1 substrates. Relative free energy surfaces for (A, C, E, G) PS1-γ-secretase and (B, D, F, H) PS2-γ-secretase in complex with substrates [denoted as substrate(cleavage position)] (A, B) NEXT(VL), (C, D) NEXT(GV), (E, F) S3(VL), (G, H) S3(AA).

The binding energies for the Notch complexes were determined by MM-GB/SA using the structures corresponding to the energetic minima derived from WTMetaD (Table 2). The data indicates that the NEXT substrate demonstrates a considerable preference for binding PS2-γ-secretase in both the Val1754/Leu1755 and Gly1753/Val1754 positions compared to PS1-γ-secretase. Similarly, the S3 substrate positioned to cleave at the Val1745/Leu1746 S4 site exhibits a preference for binding to PS2-γ-secretase. The S3 substrate positioned to cleave at the Ala1741/Ala1742 S4 site, however, preferentially binds PS1-γ-secretase.

**Table 2.** MM-GB/SA free binding energy of γ-secretase – Notch1 bound complexes.

| Substrate | Cleavage position | PS1<sup>§</sup> | PS2<sup>§</sup> |
| --- | --- | --- | --- |
| Next | V1754/L1755 | -193.1 ± 5.5 | -220.1 ± 2.1 |
| Next | G1753/V1754 | -175.6 ± 4.2 | -214.1 ± 1.6 |
| Notch-S3 | V1745/L1746 | -166.4 ± 1.1 | -199.3 ± 1.9 |
| Notch-S3 | A1741/A1742 | -181.5 ± 1.1<sup>*</sup> | -154.0 ± 2.7 |

\*The PS1-γ-secretase complex bound to Notch-S3 in the cleavage position A1741/A1742 has a second minima located at $s = 13$ to $s = 14$ and $z = 0.04$ to $z = 0.055$ , this complex conformation has a MMGBSA free binding energy of -180.4 ± 1.3 kcal/mol
<sup>§</sup>Values shown are in kcal/mol +/- standard error

### Per-residue decomposition of binding free energies for γ-secretase – Notch1 bound complexes

The preference of the NEXT substrate bound in position to cleave at the S3 site (Val1754/Leu1755) for binding to PS2-γ-secretase over PS1-γ-secretase is determined by a cluster of residues at the substrate N-terminus – Val1721 (ΔΔG_PS Pref_ = −10.29 kcal/mol), Gln1722 (ΔΔG_PS Pref_ = −2.20 kcal/mol), Ser1723 (ΔΔG_PS Pref_ = −2.91 kcal/mol) and Glu1724 (ΔΔG_PS Pref_ = −2.21 kcal/mol) – that form several hydrogen bonds and electrostatic interactions with nicastrin residue Asp655 (ΔΔG_PS Pref_ = −7.85 kcal/mol), none of which occur in the PS1-complex (Fig 7 & S4A). Further to this, the substrate residue Val1745 (ΔΔG_PS Pref_ = −2.62 kcal/mol) forms hydrophobic interactions with carbon atoms in the sidechains of PS2 Met174 (ΔΔG_PS Pref_ = −0.77 kcal/mol) and Ser175 (ΔΔG_PS Pref_ = −0.97 kcal/mol) (Fig 7 & S4B). Met174 is not conserved between PS homologues, with the analogous residue in PS1 being Ile168. The PS1 Ile168 faces outward to interact with the lipid bilayer rather than the substrate, while the PS2 Met174 interacts with the substrate. Additionally, Phe1748 (ΔΔG_PS Pref_ = −2.28 kcal/mol) in the substrate interacts with PS2 Trp171 (ΔΔG_PS Pref_ = −1.83 kcal/mol) via a π-π interaction, while PS2 Leu292 (ΔΔG_PS Pref_ = −2.27 kcal/mol) forms a CH-π bond with the substrate residue Phe1749 (ΔΔG_PS Pref_ = −1.03 kcal/mol). The protonated catalytic aspartate, PS2 Asp366 (ΔΔG_PS Pref_ = −2.76 kcal/mol), forms a hydrogen bond with the carbonyl group on the backbone of the substrate cleavage site residue, Val1754 (ΔΔG_PS Pref_ = −0.72 kcal/mol).

**Figure 7.**
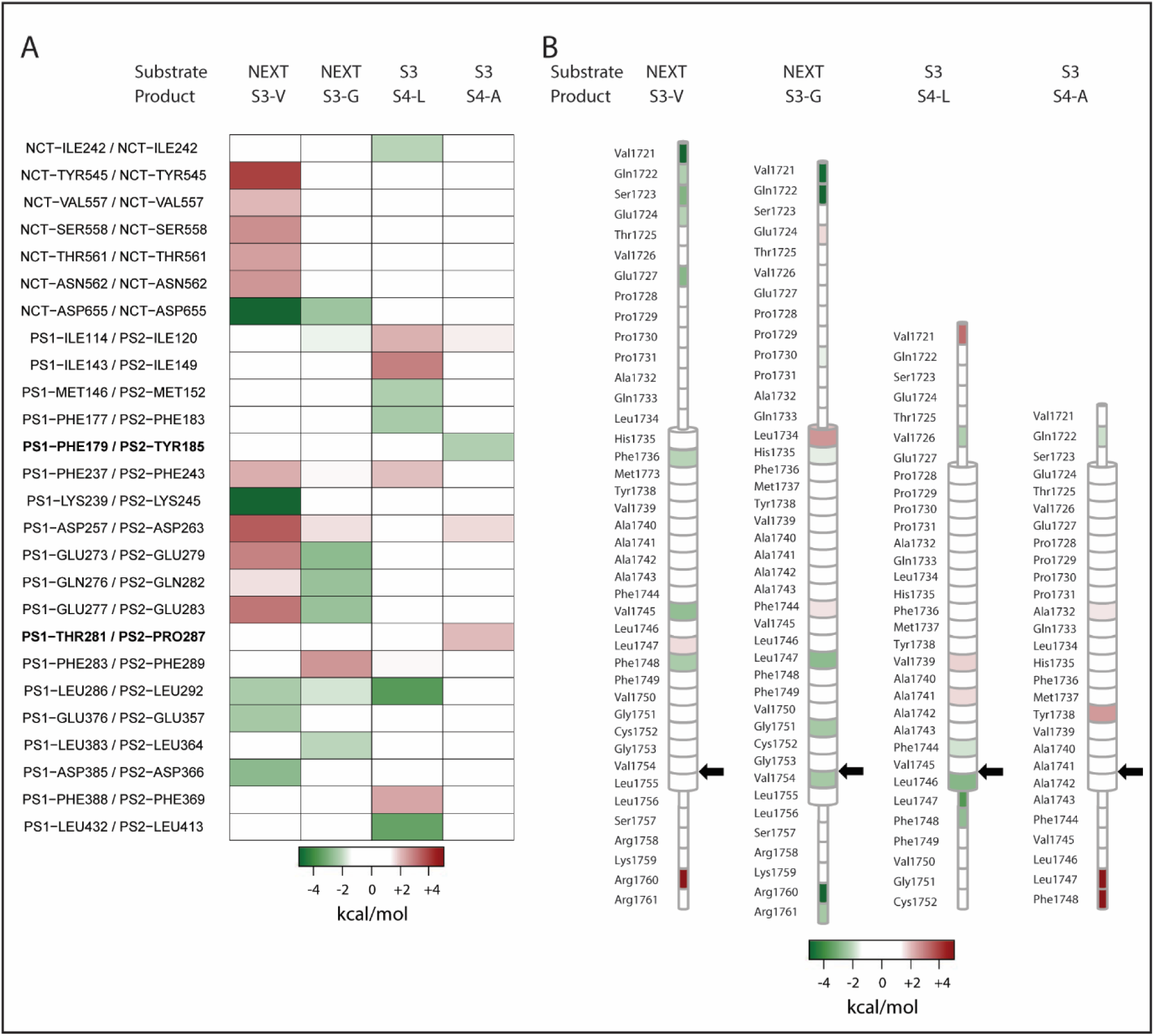
Notch1 complexes per residue heatmaps of ΔΔGPS Pref. (A) (A) Enzyme ΔΔGPS Pref values. Only residues where the ΔΔGPS Pref magnitude is greater than 1.5 kcal/mol in any complex are shown. (B) Substrate ΔΔGPS Pref values. Cleavage position denoted by arrow. Positive (red) ΔΔGPS Pref values indicate preference for PS1, negative (green) ΔΔGPS Pref values indicate preference for PS2.

Similarly, the NEXT substrate bound to be cleaved at the S3 site (Gly1753/Val1754) demonstrates a preference for PS2-γ-secretase over PS1-γ-secretase. This preference is predominantly driven by interactions with the N- and C-terminus of the substrate. At the substrate N-terminus, nicastrin residue Asp655 (ΔΔG_PS Pref_ = −2.49 kcal/mol) and the substrate residue Val1721 (ΔΔG_PS Pref_ = −4.80 kcal/mol) form a salt-bridge between the N-terminal amino group of the substrate and the aspartate side chain carboxyl group; additional hydrophobic interactions between the valine side chain and the mainchain α-carbon also occur. The adjacent substrate residue, Gln1722 (ΔΔG_PS Pref_ = −5.40 kcal/mol), forms hydrogen bonds with nicastrin residues Glu245 (ΔΔG_PS Pref_ = −0.35 kcal/mol) and Arg652 (ΔΔG_PS Pref_ = −0.67 kcal/mol) (Fig 7 & S4C). Reorganization of the substrate N-terminal loop in the PS1-complex relative to that in the PS2-complex precludes these interactions from occurring. At the substrate C-terminus, a network of hydrogen bonds and electrostatic interactions occurs between Gln282 (ΔΔG_PS Pref_ = −2.54 kcal/mol) and Glu283 (ΔΔG_PS Pref_ = −2.57 kcal/mol) with the substrate residues Arg1758 (ΔΔG_PS Pref_ = −1.27 kcal/mol), Arg1760 (ΔΔG_PS Pref_ = −6.87 kcal/mol) and Arg1761 (ΔΔG_PS Pref_ = −2.36 kcal/mol). Repositioning of the TM6a helix in the PS1-complex prevents these interactions from occurring (Fig 7 & S4D). Additional residues throughout the substrate further contribute to the preference for the PS2-complex, including: Leu1747 (ΔΔG_PS Pref_ = −2.73 kcal/mol), which forms CH-π interactions with PS2 Trp171 (ΔΔG_PS Pref_ = −0.83 kcal/mol); Gly1751 (ΔΔG_PS Pref_ = −2.08 kcal/mol), where the carbonyl group forms a hydrogen bond with the main chain amide group of PS2 Leu364 (ΔΔG_PS Pref_ = −2.05 kcal/mol); and Val1754 (ΔΔG_PS Pref_ = −2.35 kcal/mol) (the C-terminal-most residue of the substrate cleavage site), where a hydrogen bond occurs between the main chain carbonyl group and the main chain amine group of PS2 Ala415 (ΔΔG_PS Pref_ = −0.90 kcal/mol), which occurs in the PALP motif of PS2. None of these interactions are evident in the equivalent PS1-substrate complex.

The binding energies for the enzyme bound to the S3 substrate in position for the final γ-secretase cleavage at S4 site Val1745/Leu1746 suggest a substrate binding preference for PS2-γ-secretase (Table 2). This preference is supported by three key interacting regions. At the luminal juxtamembrane region, the substrate residue Val1726 (ΔΔG_PS Pref_ = −2.21 kcal/mol) forms hydrophobic interactions with nicastrin residues Phe240 and Ile242 (ΔΔG_PS Pref_ = −2.08 kcal/mol), which do not occur in the PS1-complex. Two PS2 residues, Met152 (ΔΔG_PS Pref_ = - 2.25 kcal/mol) and Phe183 (ΔΔG_PS Pref_ = −2.29 kcal/mol), co-ordinate CH-π interactions with substrate residues Tyr1738 and Met1737 respectively, that do not occur in the PS1-complex (Fig 7 & S4E). Lastly, a cluster of substrate residues around the cleavage site and PS2 residues in spatial proximity to these substrate residues contribute to the PS2-complex preference. Several polar interactions, a CH-π interaction and hydrophobic interactions are evident between substrate residues Phe1744 (ΔΔG_PS Pref_ = −1.80 kcal/mol), Leu1746 (ΔΔG_PS Pref_ = −2.86 kcal/mol), Leu1747 (ΔΔG_PS Pref_ = −3.65 kcal/mol), and Phe1748 (ΔΔG_PS Pref_ = −2.68 kcal/mol) with PS2 residues Leu292 (ΔΔG_PS Pref_ = −3.55 kcal/mol), Lys361, Gly363, Gly365 and Leu413 (ΔΔG_PS Pref_ = −3.35 kcal/mol), the majority of which do not occur in the PS1-complex (Fig 7 & S4F).

The S3 substrate in position to cleave at the Ala1741/Ala1742 S4 site for peptide release shows a preference for binding to PS1-γ-secretase over PS2-γ-secretase, unlike the other Notch complexes. This preference for binding to PS1-γ-secretase is predominantly driven by the interactions of the substrate C-terminal residues. Phe1748 (ΔΔG_PS Pref_ = +5.23 kcal/mol) and Leu1747 (ΔΔG_PS Pref_ = +6.14 kcal/mol) utilize the C-terminal carboxyl group and main chain carbonyl group, respectively, to form a salt bridge and hydrogen bonds with PS1 Lys380 (ΔΔG_PS Pref_ = +1.40 kcal/mol). Leu1747 (ΔΔG_PS Pref_ = +6.14 kcal/mol) forms hydrophobic interactions with the carbons along the length of the sidechain of Thr281 (ΔΔG_PS Pref_ = +2.03 kcal/mol) and Arg377 (ΔΔG_PS Pref_ = +1.45 kcal/mol) (Fig 7 & S4G). None of these interactions occur in the PS2 complex, and notably, the PS1 residue Thr281 is not conserved in PS2, with the analogous residue being Pro287. Further contributions to the PS1-complex preference occur within the substrate transmembrane domain, facilitated by Ala1732 (ΔΔG_PS Pref_ = +1.58 kcal/mol), which forms hydrophobic interactions with the side chains of PS1 residues Trp165 (ΔΔG_PS Pref_ = +0.67 kcal/mol), and Ile168 (ΔΔG_PS Pref_ = +0.29 kcal/mol), and Tyr1738 (ΔΔG_PS Pref_ = +2.56 kcal/mol), where the main chain carbonyl forms a hydrogen bond with the amide of PS1 Gly384 (ΔΔG_PS Pref_ = +0.29 kcal/mol). Neither interaction occurs in the PS2-complex (Fig 7 & S4H).

## DISCUSSION

The propensity for either PS1- or PS2-γ-secretase to generate a specific profile of Aβ species is a function of both the initial cleavage site and the subsequent likelihood of the successive tri/tetra-peptide cleavage events occurring. In this study, we undertook well-tempered metadynamics simulations of PS1- and PS2-γ-secretase enzymes complexes with the initial and intermediate APP substrates of the two major Aβ species, as well as Notch1-derived substrates. We analyzed these simulations to determine the likely low energy states for γ-secretase-substrate complexes and the binding free energy for each substrate bound to γ-secretase. All data was generated and compared for both PS1- and PS2-γ-secretase enzymes in order to assess enzyme preference for given substrates.

Our metadynamics results suggest a comparable ability for PS1-γ-secretase complexes to initiate either the Aβ40 or Aβ42 pathway (Fig 2A, 3A), while binding of subsequent substrates in the Aβ40 pathway (Fig 2B-D) involves a restricted conformational ensemble and may be less favorable. PS1-γ-secretase binding free energy results indicate a preference for binding APP in the position to initiate the Aβ42 pathway. Interestingly, subsequent substrates in both pathways are generally more efficiently bound, suggesting that PS1-γ-secretase processing of APP likely leads to the release of shorter Aβ peptides (Table 1). The metadynamics results for PS2-γ-secretase suggest a preference for PS2-γ-secretase to initiate the Aβ42 pathway over the Aβ40 pathway, marked by a less restricted verses a more restricted conformational ensemble for binding the respective substrates (Fig 2B, 3B). The subsequent substrates in each pathway, however, elicit broad conformational flexibility in PS2-γ-secretase, with the exception of the Aβ42(Aβ38) substrate, which yields a restricted conformational ensemble and may suggest reduced propensity for PS2-γ-secretase to generate Aβ38 products (Fig 3H). Binding free energy results for PS2-γ-secretase complexes support a considerable preference for binding APP-CTF to initiate the Aβ42 pathway. Interestingly, PS2-γ-secretase generally binds subsequent substrates in both pathways with lower binding energy (Table 1), suggesting that PS2-γ-secretase may generate longer Aβ products.

While it is likely that Notch undergoes similar tri- and tetra-peptide successive cleavage, this detail has not been elucidated; however, multiple initial cleavage (S3 sites) and final cleavage (S4 sites) γ-secretase sites have been identified.^28, 29, 56^ Here, we investigated PS1- and PS2-γ-secretase binding to Notch1 substrates aligned with the primary S3 and S4 sites. While PS1- and PS2-γ-secretase-complexes bound to the NEXT substrates in the S3 cleavage site positions elicit similar conformational flexibility in γ-secretase, the conformation of γ-secretase suggested by metadynamics simulations in the PS2 complexes is more akin to the cryoEM PS1-Notch complex (PDB 6IDF) than the PS1 complexes (Fig 6A-D). Furthermore, the PS2:NEXT(VL) and PS2:NEXT(GV) complexes have considerably more favorable binding energies over the equivalent complexes with PS1 (Table 2). Our data suggests that PS2-γ-secretase would preferentially process Notch1 substrates to generate the NICD over PS1-γ-secretase.

Generation of AICD and NICD products is indicative of the propensity for the initial cleavage by PS1- and PS2-γ-secretase, however, experimental data identifying PS1- or PS2-γ-secretaseICD generation levels is discordant. Some studies show that PS1-γ-secretase generates more AICD (or APP-CTF accumulation) or NICD, ^57–61^ while others show that PS1 and PS2 produce similar levels of ICD products.^17, 50^ Notably, these studies are undertaken in different experimental conditions, use different cell lines, and importantly, do not account for differences in PS1 and PS2 expression that will likely influence total γ-secretase activity levels. We have recently shown in HEK293 cells that PS1 expression is approximately 5-times that of PS2 expression and that this expression profile is retained in an exogenous expression system. Subsequently, we show that when PS expression is accounted for PS2-γ-secretase processes more APP and Notch substrate than PS1-γ-secretase in an exogenous system.^53^ In this study, we see that both PS1- and PS2-γ-secretase have comparable conformational flexibility when bound to NEXT substrates (Fig 6A-D). However, the substantial preference for PS2-γ-secretase binding of NEXT substrates (Table 2) shown in this study supports the notion that PS2-γ-secretase would generate more NICD product at an individual enzymatic level.

With respect to APP processing, it is the initial cleavage position and the different propensities for PS1-vs PS2-γ-secretase to cleave these that is important in the context of AD and Aβ generation. However, any preference for the initial cleavage site between PS1- and PS2-γ-secretase remains contentious; one study^62^, using PS1+/+ PS2+/+ genotype HEK293 and HeLa cells, shows that the AICD products aligning with Aβ42 pathway initiation predominates in endosomal fractions, where PS2 localises,^50, 60^ whereas AICD products aligning with Aβ40 pathway initiation predominate in plasma membrane fractions, where PS1 primarily localises.^50, 60^ Another study shows that both PS1- and PS2-γ-secretase generate similar ratios of the AICD product of both Aβ40 and Aβ42 pathways.^63^ While studies investigating the gamut of Aβ products by both PS1- and PS2-γ-secretase are not plentiful, PS2-γ-secretase has been shown to generate a higher Aβ42:Aβ40 ratio than PS1-γ-secretase.^47, 52, 53, 64^ Additionally, PS1-γ-secretase generates higher levels of Aβ38 than Aβ42, while the opposite is true of PS2-γ-secretase,^57, 59, 63, 65^ The final Aβ profile generated by γ-secretase is affected not only by the initial cleavage site, but also by the likelihood of continued processing. The data presented in this study shows that PS2-γ-secretase not only has a preference for binding to CTF(Aβ48), but also a broader conformational ensemble when binding this substrate, compared with CTF(Aβ49) (Table 1, Fig 2A-B, 3A-B), supporting the view that PS2-γ-secretase is likely the predominant Aβ42-generating enzyme. This is further supported by comparably unfavorable free binding energies of the PS2-γ-secretase bound to subsequent substrates of the Aβ42 pathway (Table 1), and the restricted conformational ensembles observed for the PS2:Aβ42(Aβ38) complex (Fig 3H). Combined, these data indicate that PS2-γ-secretase will preferentially initiate Aβ42 pathway and will likely release the substrate prior to Aβ38 product generation.

Improved understanding of PS1- and PS2-γ-secretase specific substrate processivity and the repertoire of enzyme-substrate conformations is critical for the future development of novel γ-secretase targeting therapeutics. Following the failures of γ-secretase inhibitors to gain traction, attention has turned to the development of molecules that selectively inhibit the processing of APP – γ-secretase modulators (GSM). These molecules functionally lead to increased production of shorter Aβ products, in particular Aβ38 and Aβ37, with concomitant reductions in the Aβ42 and Aβ40 products, and leave the processing of Notch and other substrate unaffected.^44, 63, 65–68^ The atomic structure of PS1-γ-secretase bound to the GSM E2012 was recently solved and from this, it has been proposed that the facilitation of substrate helix unwinding is a possible mechanism by which GSMs may increase the production of shorter Aβ products.^44^ Our study identifies conformational bottlenecks between PS1- and PS2-γ-secretase, in particular, with APP processing, presenting opportunities to stabilize or destabilize specific states. Additionally, we provide insight into γ-secretase targeting more broadly, i.e. APP vs Notch vs other substrates, as different types of substrates are shown to affect γ-secretase conformation differently, which may be harnessed for future structural based drug design.^69^ Notably, we observe different conformations between PS1- and PS2-γ-secretase bound to NEXT substrates, which implies that PS1- and PS2-complexes could be targeted differently. This is supported by PS1 vs PS2 selectivity that is already evident in γ-secretase-targeting small molecules that have been developed through traditional medicinal chemistry pipelines.^51, 59, 65, 70^ This study provides insight into the conformational repertoire of γ-secretase bound to the various substrates within a cleavage pathway, improving our understanding of substrate processivity and highlights the importance of due consideration for both PS1- and PS2-γ-secretase in structure-based drug design.

## METHODS

### Structure preparation

The structures of PS1-γ-secretase bound to APP (PDB 6IYC) and Notch1 (PDB 6IDF) were obtained from the Protein Data Bank and used for the PS1:CTF(Aβ49) and PS1:NEXT(VL) models. The subsequent PS1-γ-secretase models with either the APP or Notch substrate in different positions commensurate with the intermediate substrate and positioned for the expected product (Fig S5), and all PS2-γ-secretase models were generated using Advanced Homology Modelling within Schrodinger 2018-3. ProPKA^71^ was used to assign protonation states within prepared structures, typically predicting the catalytic aspartate Asp385/Asp366 to be protonated/neutral in charge and the catalytic aspartate Asp257/Asp263 to be deprotonated/charged. Once built, the structure was aligned to its coordinates as deposited in the Orientations of Protein in Membranes (OPM) database,^72^ to facilitate the subsequent system setup for molecular dynamics simulations. Acetyl caps at the N-terminus and N-methylamine caps at the C-terminus were added to the PS-NTF chain, while the PS-CTF chain was only capped at its N-terminus, and APH1a, NCT and the APP-CTF and NEXT substrates were only capped at the C-terminus. The PEN2 chain and subsequent substrates were not capped.

### Simulation box preparation

Built complexes were set up for simulation adapting procedures from the Amber lipid force field tutorial.^73^ Briefly, built complexes were submitted to the CHARMM-GUI web server,^74^ where they were embedded in POPC bilayers (120 x 120 lipids in size) and solvated (to create a box with a minimum distance of 15 Å between the edge of the protein and the box edge), with salt added for charge neutralization of the system and for simulating relevant physiological ionic strength (150mM NaCl). Aguayo-Ortiz et al.^39^ have shown that γ-secretase in-silico conformational behavior is similar in different homogenous membrane environments, hence we have simulated γ-secretase in 100% POPC. Relevant caps were chosen during CHARMM-GUI preparation to preserve the caps used during structure preparation.

Conversion of the prepared system from CHARMM to AMBER format, as well as system parameterization, was performed using AmberTools.^75^ The protein was parameterized using the AMBER *ff14SB* force field.^76^ Lipids were parameterized using Lipid14.^77^ TIP3P water was used throughout.^78^ The Joung-Cheatham ion parameters were used for sodium and chloride.^79^ The resulting topology was then ported to GROMACS format using *acpype*.^80^

### System Equilibration

Simulations were performed using GROMACS 2018.3^81^ patched with PLUMED 2.5.^82^ Equilibrations of the system in the NVT and NPT ensembles were adapted from previously described procedures for equilibration of membrane proteins.^83^ Briefly, heavy atoms in the protein and lipids were position-restrained with harmonic restraints at 240 kJ/(mol nm^2^), and the system gradually heated in the NVT ensemble from 0K to 100K over 0.1ns, followed by heating in the NPT ensemble from 100K to 300K for a further 0.1ns. Following this, further NPT simulations (0.1ns in duration each) were conducted with only the protein heavy atoms position-restrained, utilizing gradually decreasing force constants (120 kJ/(mol nm^2^), 96 kJ/(mol nm^2^), 72 kJ/(mol nm^2^), 48 kJ/(mol nm^2^), 24 kJ/(mol nm^2^)), with a further 0.1ns simulation performed without position restraints.

### Identification of a path between APP-bound and Notch-bound states

Targeted molecular dynamics was used to derive a path between the APP-bound and Notch1-bound states of PS1-γ-secretase. 10 simulations of 5ns duration were conducted, each starting from the APP-bound state of γ-secretase biasing towards the Notch1-bound states. During these simulations, a restraint of 50 kcal/mol was employed on the RMSD of all heavy atoms to the Notch1-bound state, as well as a concurrent added restraint of 100 kcal/mol on the RMSD of heavy atoms of presenilin transmembrane regions 2, 3 and 4, which are key interactors with the substrate. Frames from each simulation were clustered to 0.1nm, with the simulation giving rise to the largest number of clusters used to give the frames of the path. A vehicle routing solver^84^ on the RMSD matrix of the clusters was used to identify the order of clusters giving rise to the smallest distance between adjacent frames. With the exclusion of the nicastrin ectodomain, all heavy atoms were used in the final determined path.

The frames comprising the PS2-γ-secretase path were generated by homology modelling against each corresponding frame of the PS1-γ-secretase path determined by targeted molecular dynamics.

### Metadynamics Simulations of γ-Secretase complexes

After equilibration, the systems underwent well-tempered metadynamics (WTMetaD) to explore the conformational ensembles of PS1- and PS2-γ-secretase enzymes bound to APP and Notch derived substrates, similarly to that previously reported.^85^ Briefly, the collective variables for the WTMetaD bias were the position along the targeted MD-generated path (s) and the distance from this path (z). σ for s and z were set as 0.5 and 0.001 respectively, as determined by approximately half of the standard deviation in these variables at the conclusion of a 5ns unbiased simulation of PS1-γ-secretase bound to APP (Fig S6). Simulations were performed at 310 K for 500 ns, with Gaussian hills 1 kJ/mol in size added every 1 ps and employing a bias factor of 40. Calculation of the metadynamics reweighting factor^86^ was enabled and the bias was stored on a grid for computational efficiency (updated every 10 ps using grid bin widths of 0.2 in *s* and 0.0005 in *z*). Atomic coordinates, velocities and energies were saved every 10 ps. To limit the exploration of deviations from the defined path, an upper wall in *z* at 0.1 was used, with a force constant of 100 and a rescaling factor of 0.001.

### Identification of Low-Energy States of γ-Secretase complexes

Free energy surfaces (FESs) in terms of *s* and *z* were calculated from the WTMetaD simulations and reported relative to the lowest energy value in the determined FES. Convergence of FESs was assessed by monitoring the difference between free energy surfaces at 1ns intervals, as well as the Gaussian hill height and the collective variable space sampled over the duration of the simulations (Fig S7-S10). Structural ensembles for each minimum in the FES within 2.5 kJ/mol of the global minimum were extracted from the WTMetaD simulations and clustered using the GROMACS *gmx cluster* utility, employing the GROMOS algorithm^87^ and a 0.2nm threshold.

### Binding free energy calculations

The structural ensembles extracted for each energetic minimum were used to calculate the substrate-enzyme binding energies, which were determined using the molecular mechanics generalized Born/surface area (MM-GB/SA) approach, facilitated by the *MMPBSA.py* tool of AmberTools.^88^ The single-trajectory protocol was utilized,^89^ with the following equation:

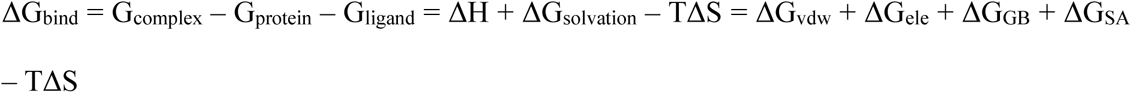

where ΔG_vdw_ is the molecular mechanics van der Waals interaction energy, ΔG_ele_ is the molecular mechanics electrostatic interaction energy, ΔG_GB_ is the change in polar desolvation energy upon complex formation and ΔG_SA_ is the change in nonpolar desolvation energy upon complex formation. The entropic term (-TΔS) was not calculated, due to high computational cost and poor accuracy. The GBneck2 model (*igb* = 5) was used to calculate the polar desolvation energy.^90^ The non-polar component of the desolvation energy was calculated via solvent accessible surface areas calculated with the LCPO method.^91^ MM-GB/SA energies were also decomposed per-residue (*idecomp* = 1).

To determine the preference between PS1- and PS2-complexes bound to the same substrate ΔΔG_PS Pref_ = ΔG_bind PS2_ - ΔG_bind PS1_ was calculated, where positive values indicate preference for PS1, while negative values indicate preference for PS2.

## AUTHOR INFORMATION

M. Eccles, G. Verdile and M. Agostino conceived this study, M. Agostino designed the experiments, M. Eccles performed the experiments and acquired the data. M. Eccles, D. Groth, G. Verdile and M. Agostino analyzed and interpreted data. All authors were involved in manuscript preparation.

## ACKNOWLEDGEMENTS

This work was supported by resources provided by the Pawsey Supercomputing Research Centre with funding from the Australian Government and the Government of Western Australia. Mark Agostino is supported by a Raine Priming Grant (Raine Medical Research Foundation) and a Curtin Research Fellowship. Melissa K Eccles is supported by a Dementia Australia Research Foundation Postgraduate Scholarship, an Australian Government Department of Education RTP Fee Offset Scholarship, and a Curtin University Postgraduate Completion Scholarship.

## ABBREVIATIONS

ADAM: A disintegrin and metalloproteinase
AD: Alzheimer’s disease
Aβ: Amyloid-β
APP: amyloid precursor protein
APH1: anterior pharynx-defective 1
AICD: APP intracellular domain
ADAD: autosomal dominant Alzheimer’s disease
BACE: β-APP cleaving enzyme
FES: free energy surface
MM-GB/SA: molecular mechanics-generalized Born/surface area
NCT: nicastrin
NICD: Notch intracellular domain
NEXT: Notch extracellular truncation
PS1: presenilin-1
PS2: presenilin-2
PEN2: presenilin enhancer 2
WTMetaD: well-tempered metadynamics
ΔΔG_bind_: binding free energy
ΔΔG_PS Pref_: difference in binding free energy between PS1 and PS2 complex

## REFERENCES

1. World Health Organisation, Towards a dementia-inclusive society: WHO toolkit for dementia-friendly initiatives (DFIs). World Health Organisation: Geneva, 2021.

2. Serrano-Pozo, A.; Frosch, M. P.; Masliah, E.; Hyman, B. T., Neuropathological alterations in Alzheimer disease. Cold Spring Harbor Perspectives in Medicine 2011, 1 (1), a006189.

3. Gravina, S. A.; Ho, L.; Eckman, C. B.; Long, K. E.; Otvos, L.; Younkin, L. H.; Suzuki, N.; Younkin, S. G., Amyloid β Protein (Aβ) in Alzheimer’s Disease Brain: Biochemical and immunocytochemical analysis with antibodies specific for forms ending at Aβ40 or Aβ42(43). Journal of Biological Chemistry 1995, 270 (13), 7013–7016.

4. Kuperstein, I.; Broersen, K.; Benilova, I.; Rozenski, J.; Jonckheere, W.; Debulpaep, M.; Vandersteen, A.; Segers-Nolten, I.; Van Der Werf, K.; Subramaniam, V.; Braeken, D.; Callewaert, G.; Bartic, C.; D’Hooge, R.; Martins, I. C.; Rousseau, F.; Schymkowitz, J.; De Strooper, B., Neurotoxicity of Alzheimer’s disease Aβ peptides is induced by small changes in the Aβ(42) to Aβ(40) ratio. The EMBO Journal 2010, 29 (19), 3408–3420.

5. Pauwels, K.; Williams, T. L.; Morris, K. L.; Jonckheere, W.; Vandersteen, A.; Kelly, G.; Schymkowitz, J.; Rousseau, F.; Pastore, A.; Serpell, L. C.; Broersen, K., Structural Basis for Increased Toxicity of Pathological Aβ42:Aβ40 Ratios in Alzheimer Disease. Journal of Biological Chemistry 2012, 287 (8), 5650–5660.

6. Scheuner, D.; Eckman, C.; Jensen, M.; Song, X.; Citron, M.; Suzuki, N.; Bird, T. D.; Hardy, J.; Hutton, M.; Kukull, W.; Larson, E.; Levy-Lahad, E.; Viitanen, M.; Peskind, E.; Poorkaj, P.; Schellenberg, G.; Tanzi, R.; Wasco, W.; Lannfelt, L.; Selkoe, D.; Younkin, S., Secreted amyloid beta-protein similar to that in the senile plaques of Alzheimer’s disease is increased in vivo by the presenilin 1 and 2 and APP mutations linked to familial Alzheimer’s disease. Nature Medicine 1996, 2 (8), 864–70.

7. Suzuki, N.; Cheung, T. T.; Cai, X. D.; Odaka, A.; Otvos, L., Jr.; Eckman, C.; Golde, T. E.; Younkin, S. G., An increased percentage of long amyloid beta protein secreted by familial amyloid beta protein precursor (beta APP717) mutants. Science 1994, 264 (5163), 1336–40.

8. Duff, K.; Eckman, C.; Zehr, C.; Yu, X.; Prada, C. M.; Perez-tur, J.; Hutton, M.; Buee, L.; Harigaya, Y.; Yager, D.; Morgan, D.; Gordon, M. N.; Holcomb, L.; Refolo, L.; Zenk, B.; Hardy, J.; Younkin, S., Increased amyloid-beta42(43) in brains of mice expressing mutant presenilin 1. Nature 1996, 383 (6602), 710–3.

9. Luo, J. E.; Li, Y.-M., Turning the tide on Alzheimer’s disease: modulation of γ-secretase. Cell & Bioscience 2022, 12 (1), 2.

10. Song, C.; Shi, J.; Zhang, P.; Zhang, Y.; Xu, J.; Zhao, L.; Zhang, R.; Wang, H.; Chen, H., Immunotherapy for Alzheimer’s disease: targeting β-amyloid and beyond. Translational Neurodegeneration 2022, 11 (1), 18.

11. Lichtenthaler, S. F.; Haass, C.; Steiner, H., Regulated intramembrane proteolysis – lessons from amyloid precursor protein processing. Journal of Neurochemistry 2011, 117 (5), 779–796.

12. Bolduc, D. M.; Montagna, D. R.; Gu, Y.; Selkoe, D. J.; Wolfe, M. S., Nicastrin functions to sterically hinder γ-secretase–substrate interactions driven by substrate transmembrane domain. Proceedings of the National Academy of Sciences of the United States of America 2016, 113 (5), E509–E518.

13. Sato, T.; Diehl, T. S.; Narayanan, S.; Funamoto, S.; Ihara, Y.; De Strooper, B.; Steiner, H.; Haass, C.; Wolfe, M. S., Active γ-secretase complexes contain only one of each component. Journal of Biological Chemistry 2007, 282 (47), 33985–33993.

14. Shah, S.; Lee, S.-F.; Tabuchi, K.; Hao, Y.-H.; Yu, C.; LaPlant, Q.; Ball, H.; Dann Iii, C. E.; Südhof, T.; Yu, G., Nicastrin Functions as a γ-Secretase-Substrate Receptor. Cell 2005, 122 (3), 435–447.

15. Wolfe, M. S.; Xia, W.; Ostaszewski, B. L.; Diehl, T. S.; Kimberly, W. T.; Selkoe, D. J., Two transmembrane aspartates in presenilin-1 required for presenilin endoproteolysis and [gamma]-secretase activity. Nature 1999, 398 (6727), 513–517.

16. Ahn, K.; Shelton, C. C.; Tian, Y.; Zhang, X.; Gilchrist, M. L.; Sisodia, S. S.; Li, Y.-M., Activation and intrinsic γ-secretase activity of presenilin 1. Proceedings of the National Academy of Sciences 2010, 107 (50), 21435–21440.

17. Yonemura, Y.; Futai, E.; Yagishita, S.; Kaether, C.; Ishiura, S., Specific combinations of presenilins and Aph1s affect the substrate specificity and activity of γ-secretase. Biochemical and Biophysical Research Communications 2016, 478 (4), 1751–1757.

18. Levy-Lahad, E.; Wasco, W.; Poorkaj, P.; Romano, D. M.; Oshima, J.; Pettingell, W. H.; Yu, C.-e.; Jondro, P. D.; Schmidt, S. D.; Wang, K.; Crowley, A. C.; Fu, Y.-H.; Guenette, S. Y.; Galas, D.; Nemens, E.; Wijsman, E. M.; Bird, T. D.; Schellenberg, G. D.; Tanzi, R. E., Candidate Gene for the Chromosome 1 Familial Alzheimer’s Disease Locus. Science 1995, 269 (5226), 973–977.

19. Sherrington, R.; Rogaev, E. I.; Liang, Y.; Rogaeva, E. A.;, et al., Cloning of a gene bearing missense mutations in early-onset familial Alzheimer’s disease. Nature 1995, 375 (6534), 754–60.

20. Tanzi, R. E.; Kovacs, D. M.; Kim, T. W.; Moir, R. D.; Guenette, S. Y.; Wasco, W., The gene defects responsible for familial Alzheimer’s disease. Neurobiology of disease 1996, 3 (3), 159–168.

21. Wu, L.; Rosa-Neto, P.; Hsiung, G.-Y. R.; Sadovnick, A. D.; Masellis, M.; Black, S. E.; Jia, J.; Gauthier, S., Early-Onset Familial Alzheimer’s Disease (EOFAD). Canadian Journal of Neurological Sciences 2012, 39 (4), 436–445.

22. Takami, M.; Nagashima, Y.; Sano, Y.; Ishihara, S.; Morishima-Kawashima, M.; Funamoto, S.; Ihara, Y., γ-Secretase: Successive tripeptide and tetrapeptide release from the transmembrane domain of β-carboxyl terminal fragment. The Journal of Neuroscience 2009, 29 (41), 13042–13052.

23. Matsumura, N.; Takami, M.; Okochi, M.; Wada-Kakuda, S.; Fujiwara, H.; Tagami, S.; Funamoto, S.; Ihara, Y.; Morishima-Kawashima, M., γ-Secretase associated with lipid rafts: multiple interactive pathways in the stepwise processing of β-carboxyl-terminal fragment. Journal of Biological Chemistry 2014, 289 (8), 5109–5121.

24. Siebel, C.; Lendahl, U., Notch Signaling in Development, Tissue Homeostasis, and Disease. Physiological Reviews 2017, 97 (4), 1235–1294.

25. Mumm, J. S.; Schroeter, E. H.; Saxena, M. T.; Griesemer, A.; Tian, X.; Pan, D. J.; Ray, W. J.; Kopan, R., A ligand-induced extracellular cleavage regulates γ-secretase-like proteolytic activation of notch1. Molecular Cell 2000, 5 (2), 197–206.

26. van Tetering, G.; van Diest, P.; Verlaan, I.; van der Wall, E.; Kopan, R.; Vooijs, M., Metalloprotease ADAM10 is required for Notch1 site 2 cleavage. Journal of Biological Chemistry 2009, 284 (45), 31018–27.

27. Schroeter, E. H.; Kisslinger, J. A.; Kopan, R., Notch-1 signalling requires ligand-induced proteolytic release of intracellular domain. Nature 1998, 393 (6683), 382–6.

28. Tagami, S.; Okochi, M.; Yanagida, K.; Ikuta, A.; Fukumori, A.; Matsumoto, N.; Ishizuka-Katsura, Y.; Nakayama, T.; Itoh, N.; Jiang, J.; Nishitomi, K.; Kamino, K.; Morihara, T.; Hashimoto, R.; Tanaka, T.; Kudo, T.; Chiba, S.; Takeda, M., Regulation of Notch signaling by dynamic changes in the precision of S3 cleavage of Notch-1. Molecular and Cellular Biology 2008, 28 (1), 165–176.

29. Okochi, M.; Steiner, H.; Fukumori, A.; Tanii, H.; Tomita, T.; Tanaka, T.; Iwatsubo, T.; Kudo, T.; Takeda, M.; Haass, C., Presenilins mediate a dual intramembranous gamma-secretase cleavage of Notch-1. The EMBO Journal 2002, 21 (20), 5408–5416.

30. Bai, X.-c.; Yan, C.; Yang, G.; Lu, P.; Ma, D.; Sun, L.; Zhou, R.; Scheres, S. H. W.; Shi, Y., An atomic structure of human γ-secretase. Nature 2015, 525 (7568), 212–217.

31. Bai, X.-c.; Rajendra, E.; Yang, G.; Shi, Y.; Scheres, S. H. W., Sampling the conformational space of the catalytic subunit of human γ-secretase. eLife 2015, 4, e11182.

32. Zhou, R.; Yang, G.; Guo, X.; Zhou, Q.; Lei, J.; Shi, Y., Recognition of the amyloid precursor protein by human γ-secretase. Science 2019, eaaw0930.

33. Yang, G.; Zhou, R.; Zhou, Q.; Guo, X.; Yan, C.; Ke, M.; Lei, J.; Shi, Y., Structural basis of Notch recognition by human γ-secretase. Nature 2018.

34. Hitzenberger, M.; Zacharias, M., Structural Modeling of γ-Secretase Aβn Complex Formation and Substrate Processing. ACS Chemical Neuroscience 2019, 10 (3), 1826–1840.

35. Somavarapu, A. K.; Kepp, K. P., Membrane Dynamics of γ-Secretase Provides a Molecular Basis for β-Amyloid Binding and Processing. ACS Chemical Neuroscience 2017.

36. Aguayo-Ortiz, R.; Chavez-Garcia, C.; Straub, J. E.; Dominguez, L., Characterizing the structural ensemble of [gamma]-secretase using a multiscale molecular dynamics approach. Chemical Science 2017, 8 (8), 5576–5584.

37. Hitzenberger, M.; Zacharias, M., γ-Secretase Studied by Atomistic Molecular Dynamics Simulations: Global Dynamics, Enzyme Activation, Water Distribution and Lipid Binding. Frontiers in Chemistry 2018, 6, 640.

38. Dehury, B.; Tang, N.; Kepp, K. P., Molecular dynamics of C99-bound γ-secretase reveal two binding modes with distinct compactness, stability, and active-site retention: implications for Aβ production. Biochemical Journal 2019, 476 (7), 1173–1189.

39. Aguayo-Ortiz, R.; Straub, J. E.; Dominguez, L., Influence of membrane lipid composition on the structure and activity of γ-secretase. Physical Chemistry Chemical Physics 2018, 20 (43), 27294–27304.

40. Chen, S.-Y.; Zacharias, M., How Mutations Perturb γ-Secretase Active Site Studied by Free Energy Simulations. ACS Chemical Neuroscience 2020, 11 (20), 3321–3332.

41. Dehury, B.; Somavarapu, A. K.; Kepp, K. P., A computer-simulated mechanism of familial Alzheimer’s disease: Mutations enhance thermal dynamics and favor looser substrate-binding to γ-secretase. Journal of Structural Biology 2020, 212 (3), 107648.

42. Mehra, R.; Kepp, K. P., Computational prediction and molecular mechanism of γ-secretase modulators. European Journal of Pharmaceutical Sciences 2021, 157, 105626.

43. Hitzenberger, M.; Zacharias, M., Uncovering the Binding Mode of γ -Secretase Inhibitors. ACS Chemical Neuroscience 2019, 10 (8), 3398–3403.

44. Yang, G.; Zhou, R.; Guo, X.; Yan, C.; Lei, J.; Shi, Y., Structural basis of γ-secretase inhibition and modulation by small molecule drugs. Cell 2021, 184 (2), 521–533.e14.

45. Landrum, M. J.; Lee, J. M.; Benson, M.; Brown, G. R.; Chao, C.; Chitipiralla, S.; Gu, B.; Hart, J.; Hoffman, D.; Jang, W.; Karapetyan, K.; Katz, K.; Liu, C.; Maddipatla, Z.; Malheiro, A.; McDaniel, K.; Ovetsky, M.; Riley, G.; Zhou, G.; Holmes, J. B.; Kattman, B. L.; Maglott, D. R., ClinVar: improving access to variant interpretations and supporting evidence. Nucleic Acids Research 2018, 46 (D1), D1062–d1067.

46. Culvenor, J. G.; Evin, G.; Cooney, M. A.; Wardan, H.; Sharples, R. A.; Maher, F.; Reed, G.; Diehlmann, A.; Weidemann, A.; Beyreuther, K.; Masters, C. L., Presenilin 2 expression in neuronal cells: induction during differentiation of embryonic carcinoma cells. Experimental Cell Research 2000, 255 (2), 192–206.

47. Watanabe, H.; Imaizumi, K.; Cai, T.; Zhou, Z.; Tomita, T.; Okano, H., Flexible and accurate substrate processing with distinct presenilin/γ-secretases in human cortical neurons. eNeuro 2021, 8 (2), ENEURO.0500–20.2021.

48. Lee, M. K.; Slunt, H. H.; Martin, L. J.; Thinakaran, G.; Kim, G.; Gandy, S. E.; Seeger, M.; Koo, E.; Price, D. L.; Sisodia, S. S., Expression of presenilin 1 and 2 (PS1 and PS2) in human and murine tissues. The Journal of Neuroscience 1996, 16 (23), 7513–7525.

49. Kumar, A.; Thakur, M. K., Presenilin 1 and 2 are expressed differentially in the cerebral cortex of mice during development. Neurochemistry International 2012, 61 (5), 778–782.

50. Sannerud, R.; Esselens, C.; Ejsmont, P.; Mattera, R.; Rochin, L.; Tharkeshwar, Arun K.; De Baets, G.; De Wever, V.; Habets, R.; Baert, V.; Vermeire, W.; Michiels, C.; Groot, Arjan J.; Wouters, R.; Dillen, K.; Vints, K.; Baatsen, P.; Munck, S.; Derua, R.; Waelkens, E.; Basi, Guriqbal S.; Mercken, M.; Vooijs, M.; Bollen, M.; Schymkowitz, J.; Rousseau, F.; Bonifacino, Juan S.; Van Niel, G.; De Strooper, B.; Annaert, W., Restricted location of PSEN2/γ-secretase determines substrate specificity and generates an intracellular Aβ pool. Cell 2016, 166 (1), 193–208.

51. Lessard, C. B.; Rodriguez, E.; Ladd, T. B.; Minter, L. M.; Osborne, B. A.; Miele, L.; Golde, T. E.; Ran, Y., Individual and combined presenilin 1 and 2 knockouts reveal that both have highly overlapping functions in HEK293T cells. Journal of Biological Chemistry 2019, 294 (29), 11276–11285.

52. Pimenova, A. A.; Goate, A. M., Novel presenilin 1 and 2 double knock-out cell line for in vitro validation of PSEN1 and PSEN2 mutations. Neurobiology of Disease 2020, 104785.

53. Eccles, M. K.; Main, N.; Sabale, M.; Roberts-Mok, B.; Agostino, M.; Groth, D.; Fraser, P. E.; Verdile, G., Quantitative Comparison of Presenilin Protein Expression Reveals Greater Activity of PS2-γ-Secretase. bioRxiv 2023, 2023.05.09.540102.

54. Barrett, P. J.; Song, Y.; Van Horn, W. D.; Hustedt, E. J.; Schafer, J. M.; Hadziselimovic, A.; Beel, A. J.; Sanders, C. R., The Amyloid Precursor Protein has a Flexible Transmembrane Domain and Binds Cholesterol. Science 2012, 336 (6085), 1168–1171.

55. Tsuzuki, S.; Honda, K.; Uchimaru, T.; Mikami, M.; Tanabe, K., Origin of the Attraction and Directionality of the NH/π Interaction: Comparison with OH/π and CH/π Interactions. Journal of the American Chemical Society 2000, 122 (46), 11450–11458.

56. Okochi, M.; Fukumori, A.; Jiang, J.; Itoh, N.; Kimura, R.; Tagami, S.; Takeda, M.; Steiner, H.; Haass, C., Secretion of the Notch-1 Aβ-like peptide during Notch signaling. Journal of Biological Chemistry 2006, 281 (12), 7890–7898.

57. Acx, H.; Chávez-Gutiérrez, L.; Serneels, L.; Lismont, S.; Benurwar, M.; Elad, N.; De Strooper, B., Signature amyloid β profiles are produced by different γ-secretase complexes. Journal of Biological Chemistry 2014, 289 (7), 4346–4355.

58. Frånberg, J.; Svensson, A. I.; Winblad, B.; Karlström, H.; Frykman, S., Minor contribution of presenilin 2 for γ-secretase activity in mouse embryonic fibroblasts and adult mouse brain. Biochemical and Biophysical Research Communications 2011, 404 (1), 564–568.

59. Lee, J.; Song, L.; Terracina, G.; Bara, T.; Josien, H.; Asberom, T.; Sasikumar, T. K.; Burnett, D. A.; Clader, J.; Parker, E. M.; Zhang, L., Identification of presenilin 1-selective γ-secretase inhibitors with reconstituted γ-secretase complexes. Biochemistry 2011, 50 (22), 4973–4980.

60. Meckler, X.; Checler, F., Presenilin 1 and presenilin 2 target γ-secretase complexes to distinct cellular compartments. Journal of Biological Chemistry 2016, 291 (24), 12821–12837.

61. Zhang, Z.; Nadeau, P.; Song, W.; Donoviel, D.; Yuan, M.; Bernstein, A.; Yankner, B. A., Presenilins are required for [gamma]-secretase cleavage of [beta]-APP and transmembrane cleavage of Notch-1. Nature Cell Biology 2000, 2 (7), 463–5.

62. Fukumori, A.; Okochi, M.; Tagami, S.; Jiang, J.; Itoh, N.; Nakayama, T.; Yanagida, K.; Ishizuka-Katsura, Y.; Morihara, T.; Kamino, K.; Tanaka, T.; Kudo, T.; Tanii, H.; Ikuta, A.; Haass, C.; Takeda, M., Presenilin-Dependent γ-Secretase on Plasma Membrane and Endosomes Is Functionally Distinct. Biochemistry 2006, 45 (15), 4907–4914.

63. Lessard, C. B.; Rodriguez, E.; Ladd, T. B.; Minter, L. M.; Osborne, B. A.; Miele, L.; Golde, T. E.; Ran, Y., γ-Secretase modulators exhibit selectivity for modulation of APP cleavage but inverse γ-secretase modulators do not. Alzheimer’s Research & Therapy 2020, 12 (1), 61.

64. Placanica, L.; Tarassishin, L.; Yang, G.; Peethumnongsin, E.; Kim, S.-H.; Zheng, H.; Sisodia, S. S.; Li, Y.-M., Pen2 and presenilin-1 modulate the dynamic equilibrium of presenilin-1 and presenilin-2 γ-secretase complexes. Journal of Biological Chemistry 2009, 284 (5), 2967–2977.

65. Ebke, A.; Luebbers, T.; Fukumori, A.; Shirotani, K.; Haass, C.; Baumann, K.; Steiner, H., Novel γ-secretase enzyme modulators directly target presenilin protein. Journal of Biological Chemistry 2011, 286 (43), 37181–6.

66. Kounnas, M. Z.; Danks, A. M.; Cheng, S.; Tyree, C.; Ackerman, E.; Zhang, X.; Ahn, K.; Nguyen, P.; Comer, D.; Mao, L.; Yu, C.; Pleynet, D.; Digregorio, P. J.; Velicelebi, G.; Stauderman, K. A.; Comer, W. T.; Mobley, W. C.; Li, Y.-M.; Sisodia, S. S.; Tanzi, R. E.; Wagner, S. L., Modulation of γ-Secretase Reduces β-Amyloid Deposition in a Transgenic Mouse Model of Alzheimer’s Disease. Neuron 2010, 67 (5), 769–780.

67. Pozdnyakov, N.; Murrey, H. E.; Crump, C. J.; Pettersson, M.; Ballard, T. E.; am Ende, C. W.; Ahn, K.; Li, Y.-M.; Bales, K. R.; Johnson, D. S., γ-Secretase Modulator (GSM) Photoaffinity Probes Reveal Distinct Allosteric Binding Sites on Presenilin. Journal of Biological Chemistry 2013, 288 (14), 9710–9720.

68. Kounnas, M. Z.; Lane-Donovan, C.; Nowakowski, D. W.; Herz, J.; Comer, W. T., NGP 555, a γ-secretase modulator, lowers the amyloid biomarker, Aβ42, in cerebrospinal fluid while preventing Alzheimer’s disease cognitive decline in rodents. Alzheimer’s & Dementia: Translational Research & Clinical Interventions 2017, 3 (1), 65–73.

69. Ioppolo, A.; Eccles, M.; Groth, D.; Verdile, G.; Agostino, M., Evaluation of Virtual Screening Strategies for the Identification of γ-Secretase Inhibitors and Modulators. Molecules 2021, 27 (1).

70. Guo, X.; Wang, Y.; Zhou, J.; Jin, C.; Wang, J.; Jia, B.; Jing, D.; Yan, C.; Lei, J.; Zhou, R.; Shi, Y., Molecular basis for isoform-selective inhibition of presenilin-1 by MRK-560. Nature Communications 2022, 13 (1), 6299.

71. Madhavi Sastry, G.; Adzhigirey, M.; Day, T.; Annabhimoju, R.; Sherman, W., Protein and ligand preparation: parameters, protocols, and influence on virtual screening enrichments. Journal of Computer-Aided Molecular Design 2013, 27 (3), 221–234.

72. Lomize, M. A.; Pogozheva, I. D.; Joo, H.; Mosberg, H. I.; Lomize, A. L., OPM database and PPM web server: resources for positioning of proteins in membranes. Nucleic Acids Research 2012, 40 (Database issue), D370–D376.

73. Madej, B. D.; Walker, R. C. An Amber Lipid Force Field Tutorial: Lipid14 Edition. http://ambermd.org/tutorials/advanced/tutorial16/ (accessed 30th October 2018).

74. Jo, S.; Cheng, X.; Lee, J.; Kim, S.; Park, S.-J.; Patel, D. S.; Beaven, A. H.; Lee, K. I.; Rui, H.; Roux, B.; MacKerell, A. D.; Klauda, J. B.; Qi, Y.; Im, W., CHARMM-GUI 10 Years for Biomolecular Modeling and Simulation. Journal of Computational Chemistry 2017, 38 (15), 1114–1124.

75. Salomon-Ferrer, R.; Case, D. A.; Walker, R. C., An overview of the Amber biomolecular simulation package. Wiley Interdisciplinary Reviews: Computational Molecular Science 2013, 3 (2), 198–210.

76. Maier, J. A.; Martinez, C.; Kasavajhala, K.; Wickstrom, L.; Hauser, K. E.; Simmerling, C., ff14SB: Improving the Accuracy of Protein Side Chain and Backbone Parameters from ff99SB. Journal of Chemical Theory and Computation 2015, 11 (8), 3696–3713.

77. Dickson, C. J.; Madej, B. D.; Skjevik, Å. A.; Betz, R. M.; Teigen, K.; Gould, I. R.; Walker, R. C., Lipid14: The Amber Lipid Force Field. Journal of Chemical Theory and Computation 2014, 10 (2), 865–879.

78. Jorgensen, W. L.; Chandrasekhar, J.; Madura, J. D., Comparison of simple potential functions for simulating liquid water. The Journal of Chemical Physics 1983, 79 (2), 926–935.

79. Joung, I. S.; Cheatham, T. E., Determination of Alkali and Halide Monovalent Ion Parameters for Use in Explicitly Solvated Biomolecular Simulations. The Journal of Physical Chemistry B 2008, 112 (30), 9020–9041.

80. Sousa da Silva, A. W.; Vranken, W. F., ACPYPE - AnteChamber PYthon Parser interfacE. BMC Research Notes 2012, 5 (1), 367.

81. Abraham, M. J.; Murtola, T.; Schulz, R.; Páll, S.; Smith, J. C.; Hess, B.; Lindahl, E., GROMACS: High performance molecular simulations through multi-level parallelism from laptops to supercomputers. SoftwareX 2015, 1-2, 19–25.

82. Tribello, G. A.; Bonomi, M.; Branduardi, D.; Camilloni, C.; Bussi, G., PLUMED 2: New feathers for an old bird. Computer Physics Communications 2014, 185 (2), 604–613.

83. Huang, W.; Manglik, A.; Venkatakrishnan, A. J.; Laeremans, T.; Feinberg, E. N.; Sanborn, A. L.; Kato, H. E.; Livingston, K. E.; Thorsen, T. S.; Kling, R.; Granier, S.; Gmeiner, P.; Husbands, S. M.; Traynor, J. R.; Weis, W. I.; Steyaert, J.; Dror, R. O.; Kobilka, B. K., Structural insights into μ-opioid receptor activation. Nature 2015, 524 (7565), 315–321.

84. Perron, L.; Furnon, V., OR-Tools v7.2. Google.

85. Agostino, M.; McKenzie, F.; Buck, C.; Woodward, K. J.; Atkinson, V. J.; Azmanov, D. N.; Heng, J. I.-T., Studying Disease-Associated UBE3A Missense Variants Using Enhanced Sampling Molecular Simulations. ACS Omega 2022, 7 (29), 25039–25045.

86. Tiwary, P.; Parrinello, M., A time-independent free energy estimator for metadynamics. The Journal of Physical Chemistry B 2015, 119 (3), 736–742.

87. Daura, X.; Gademann, K.; Jaun, B.; Seebach, D.; Van Gunsteren, W. F.; Mark, A. E., Peptide folding: when simulation meets experiment. Angewandte Chemie International Edition 1999, 38 (1-2), 236–240.

88. Miller, B. R.; McGee, T. D.; Swails, J. M.; Homeyer, N.; Gohlke, H.; Roitberg, A. E., MMPBSA.py: An Efficient Program for End-State Free Energy Calculations. Journal of Chemical Theory and Computation 2012, 8 (9), 3314–3321.

89. Gohlke, H.; Kiel, C.; Case, D. A., Insights into Protein–Protein Binding by Binding Free Energy Calculation and Free Energy Decomposition for the Ras–Raf and Ras–RalGDS Complexes. Journal of Molecular Biology 2003, 330 (4), 891–913.

90. Nguyen, H.; Roe, D. R.; Simmerling, C., Improved Generalized Born Solvent Model Parameters for Protein Simulations. Journal of Chemical Theory and Computation 2013, 9 (4), 2020–2034.

91. Jörg, W.; S., S.P.; Clark, S. W., Approximate atomic surfaces from linear combinations of pairwise overlaps (LCPO). Journal of Computational Chemistry 1999, 20 (2), 217–230.

