## Supplemental Data for "Presenilin homologues influence substrate binding and processing by γ-secretase: a molecular simulation study"

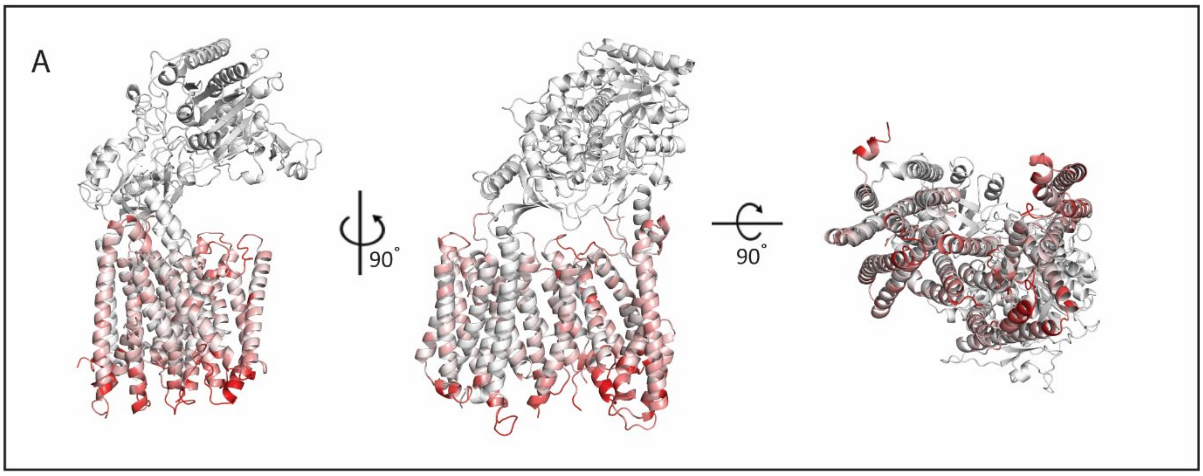


**Figure S1. Path derivation between APP and Notch1 bound states.** (A) Root Mean Squared Fluctuation (RMSF) range of 0.00nm (white) to 0.20nm (red) of PS1, Aph1a, and Pen-2 components used for path derivation.

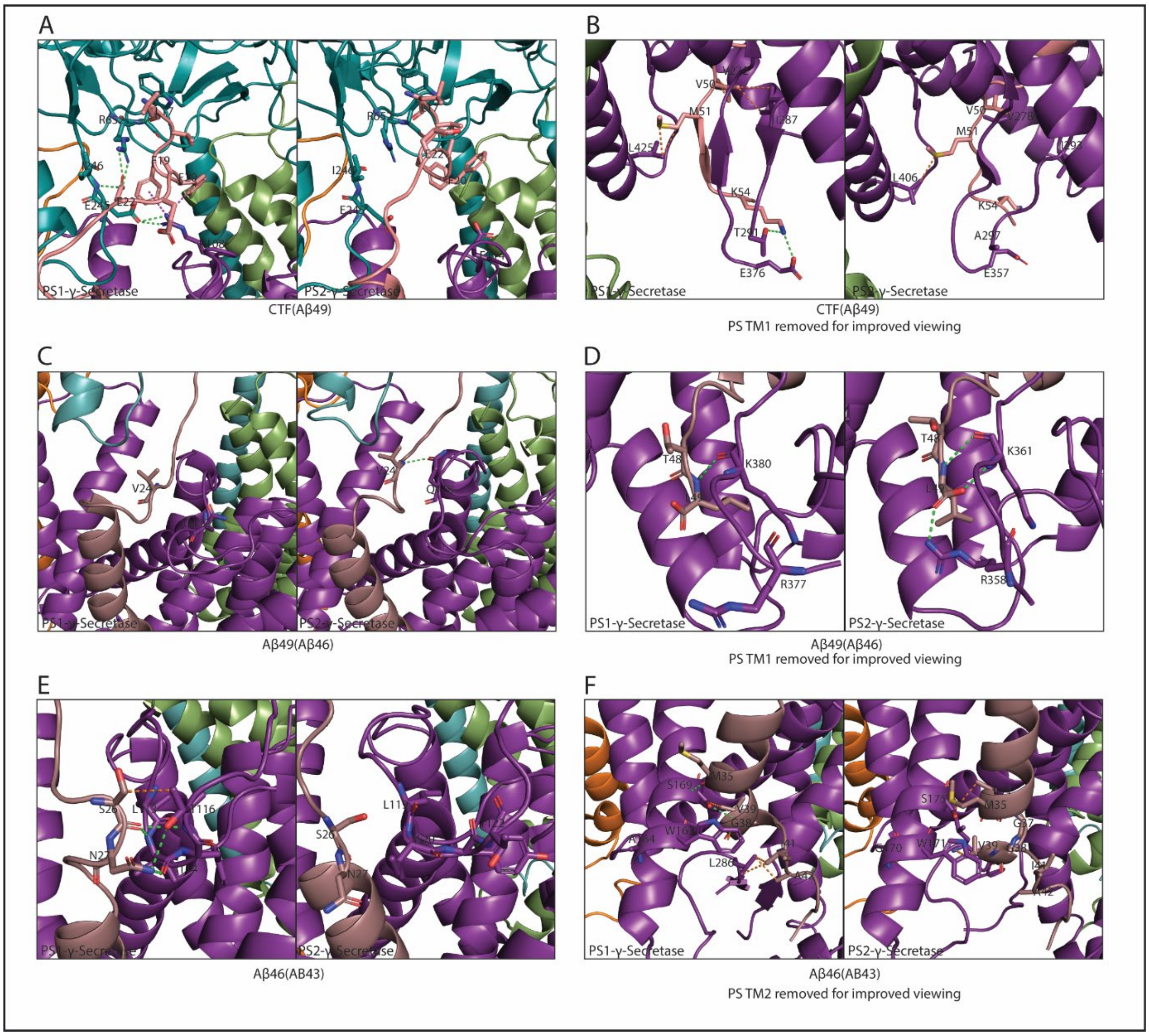


**Figure S2. Representative structures of substrate – enzyme molecular interactions contributing to $\Delta\Delta G_{PS\ Pref}$ in Aβ40 pathway.** (A) APP-CTF(Aβ49) substrate N-terminal residues Leu17 to Glu22 (B) APP-CTF(Aβ49) substrate C-terminal residues Val50 to Lys54 (C) Aβ49(Aβ46) substrate N-terminal juxta-membrane residues Val24 (D) Aβ49(Aβ46) substrate N-terminal residues Thr48 – Leu49 (E) Aβ46(Aβ43) substrate N-terminal juxta-membrane Ser26 – Asn27 (F) Aβ46(Aβ43) substrate C-terminal inner leaflet – juxta-membrane residues Met35 – Ala42. Complex components represented in cartoon coloured as nicastrin = teal, presenilin1/presenilin2 = purple, Aph1 = green, Pen-2 = orange, substrate = light pink, with specific residues involved in interactions depicted in stick format. Hydrogen bonds represented in green dashed lines, $\pi$ interactions represented by purple dashed lines, hydrophobic interactions represented in orange dashed lines.

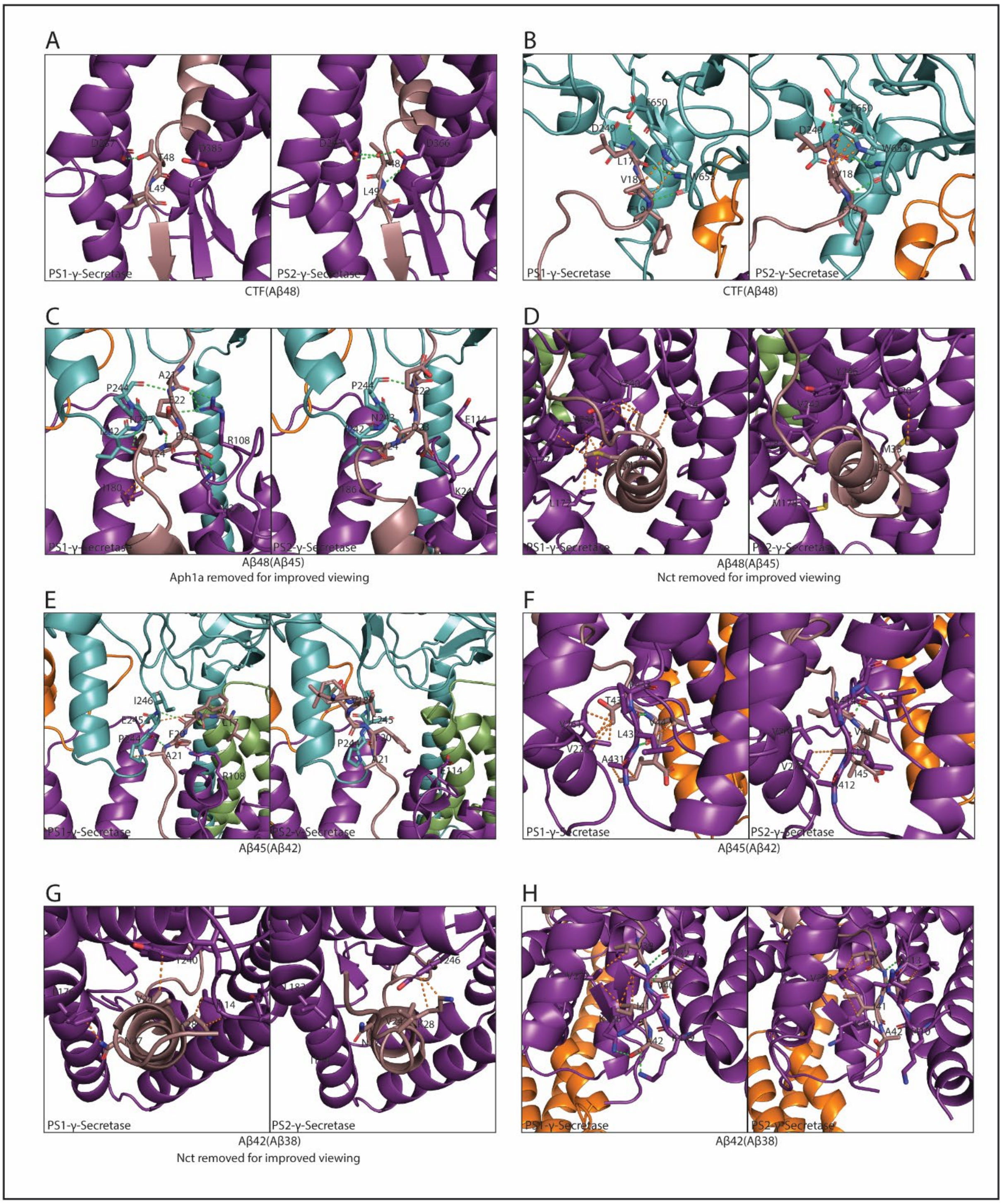


**Figure S3. Representative structures of substrate – enzyme molecular interactions contributing to ΔΔG$_{PS\ Pref}$ in Aβ42 pathway.** (A) APP-CTF(Aβ48) substrate cleavage site residues Thr48 – Leu49 (B) APP-CTF(Aβ48) substrate N-terminal residues Leu17 – phe19 (C) Aβ48(Aβ45) substrate N-terminal juxta-membrane residues Ala21 – Val24 (D) Aβ48(Aβ45) substrate TMD residues Ile32 – Met35 (E) Aβ45(Aβ42) substrate N-terminal residues Leu17 – Ala21 (F) Aβ45(Aβ42) substrate C-terminal residues Thr43 – Ile45 (G) Aβ42(Aβ38) N-terminal juxta-membrane residues Val24 – Lys28 (H) Aβ42(Aβ38) C-terminal residues Val39 – Ala42. Complex components represented in cartoon coloured as nicastrin = teal, presenilin1/presenilin2 = purple, Aph1 = green, Pen-2 = orange, substrate = light pink, with specific residues involved in interactions depicted in stick format. Hydrogen bonds represented in green dashed lines, π interactions represented by purple dashed lines, hydrophobic interactions represented in orange dashed lines.

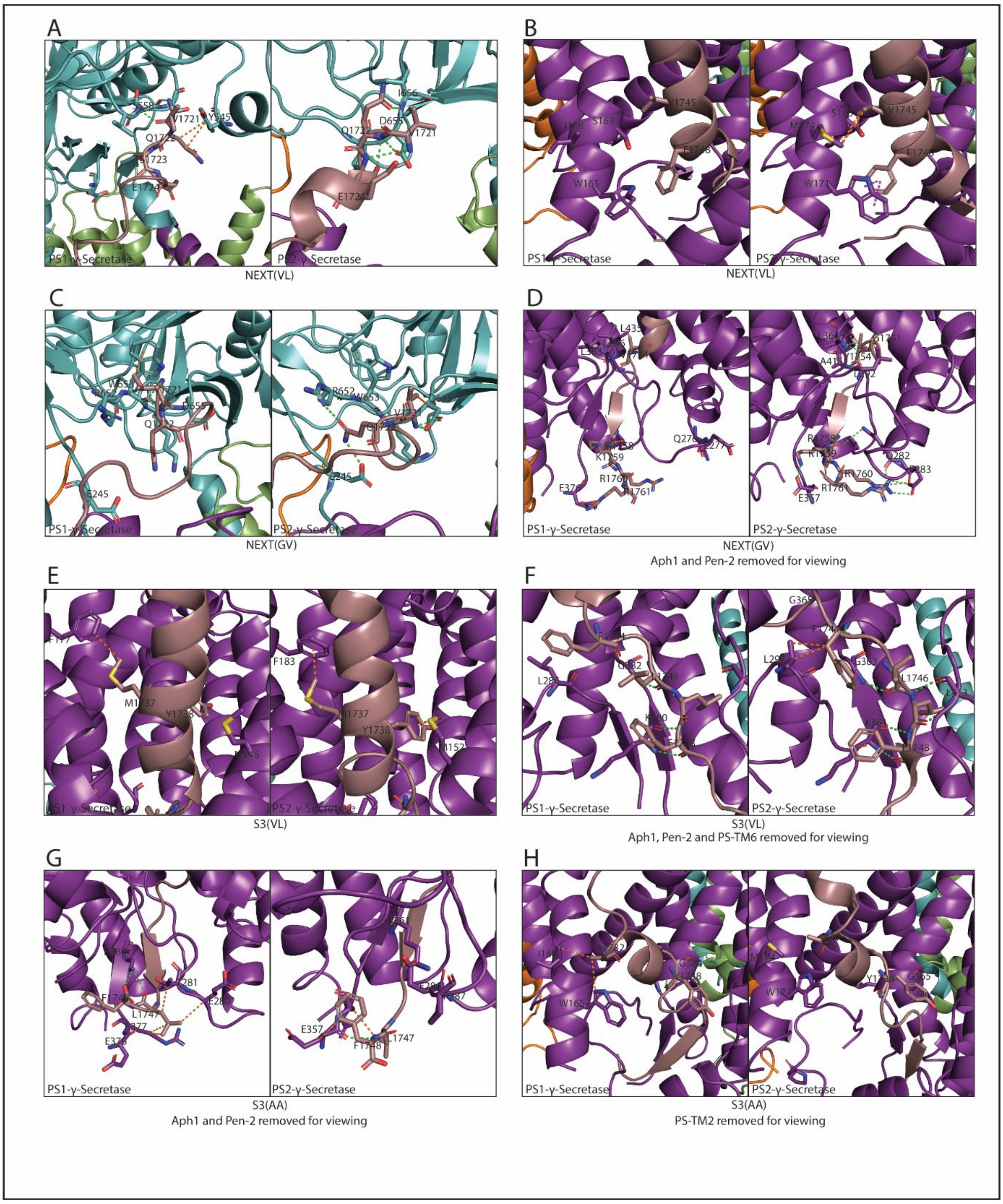


**Figure S4. Representative structures of substrate – enzyme molecular interactions contributing to $\Delta\Delta G_{PS\ Pref}$ in Notch1 cleavage.** (A) NEXT(VL) substrate N-terminus residues Val1721 – Glu1724 (B) NEXT(VL) substrate TMD residues Val1745 – Phe1748 (C) NEXT(GV) substrate N-terminal residues Val1721 – Gln1722 (D) NEXT(GV) substrate C-terminal inner leaflet – juxta-membrane residues Gly1751 – Arg1761 (E) S3(VL) substrate

TMD residues Met1737 – Tyr1738 (F) S3(VL) substrate C-terminal juxta-membrane residues Phe1744 – Phe1748 (G) S3(AA) substrate C-terminal residues Leu1747 – Phe1748 (H) S3(AA) substrate TMD residues Ala1732 – Tyr1738. Complex components represented in cartoon coloured as nicastrin = teal, presenilin1/presenilin2 = purple, Aph1 = green, Pen-2 = orange, substrate = light pink, with specific residues involved in interactions depicted in stick format. Hydrogen bonds represented in green dashed lines, π interactions represented by purple dashed lines, hydrophobic interactions represented in orange dashed lines.

```
A
    6IYC                 LVFFAEDVGSNKGAIIGLMVGGVVIATVIVITLVMLKKK
    APP-CTF - Aβ49       LVFFAEDVGSNKGAIIGLMVGGVVIATVIVITLVMLKKK

    6IYC                 LVFFAEDVGSNKGAIIGLMVGGVVIATVIVITLVMLKKK
    Aβ49 - Aβ46          LVFFAEDV---GSNKGAIIGLMVGGVVIATVIVITL

    6IYC                 LVFFAEDVGSNKGAIIGLMVGGVVIATVIVITLVMLKKK
    Aβ46 - Aβ43          LVFFAE------DVGSNKGAIIGLMVGGVVIATVIV

    6IYC                 LVFFAEDVGSNKGAIIGLMVGGVVIATVIVITLVMLKKK
    Aβ43 - Aβ40          ---LV------FFAEDVGSNKGAIIGLMVGGVVIAT

B
    6IYC                 LVFFAEDVGSNKGAIIGLMVGGVVIATVIVITLVMLKKK
    APP-CTF - Aβ48       LVFFAEDV-GSNKGAIIGLMVGGVVIATVIVITLVMLKKK

    6IYC                 LVFFAEDVGSNKGAIIGLMVGGVVIATVIVITLVMLKKK
    Aβ48 - Aβ45          LVFFAE----DVGSNKGAIIGLMVGGVVIATVIVIT

    6IYC                 LVFFAEDVGSNKGAIIGLMVGGVVIATVIVITLVMLKKK
    Aβ45 - Aβ42          -LVFFA------EDVGSNKGAIIGLMVGGVVIATVI

    6IYC                 LVFFAEDVGSNKGAIIGLMVGGVVIATVIVITLVMLKKK
    Aβ42 - Aβ38          -----------LVFFAEDVGSNKGAIIGLMVGGVVIA

C
    6IDF                 VQSETVEPPPPAQLHFMYVAAAAFVLLFFVGCGVLLSRKRR
    NEXT - S3VL          VQSETVEPPPPAQLHFMYVAAAAFVLLFFVGCGVLLSRKRR

    6IDF                 VQSETVEP-PPPAQLHFMYVAAAAFVLLFFVGCGVL-LSRKRR
    NEXT - S3GV          -VQSETVEPPPPA-QLHFMYVAAAAFVLLFFVGCGVLLSRKR

    6IDF                 VQSETVEPPPPAQLHFMYVAAAAFVLLFFVGCGVLLSRKRR
    S3GV - S4VL          ---------VQSETVEPPPPAQLHFMYVAAAAFVLLFFVGC

    6IDF                 VQSETVEPPPPAQLHFMYVAAAAFVLLFFVGCGVLLSRKRR
    S3GV - S4AA          -------------VQSETVEPPPPAQLHFMYVAAAAFVLLF
```

**Figure S5. Sequence alignments used for homology modelling of substrates**. (A) Aβ40 pathway, (B) Aβ42 pathway and (C) Notch1 cleavage.

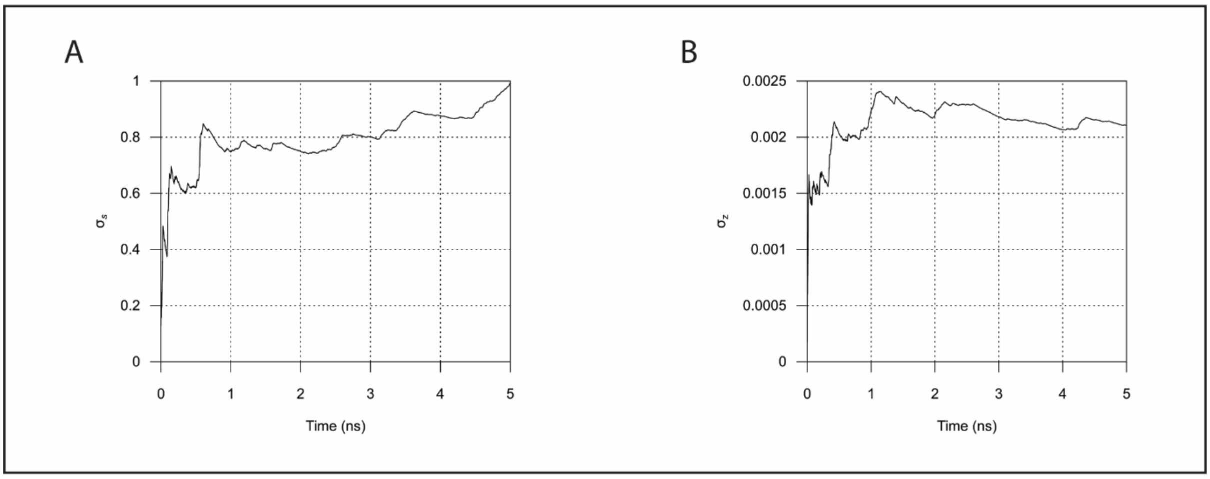


**Figure S6. Determination of σ for s and z**. (A) $\sigma_s$ and (B) $\sigma_Z$ from 5ns unbiased simulation of PS1-γ-secretase bound to APP

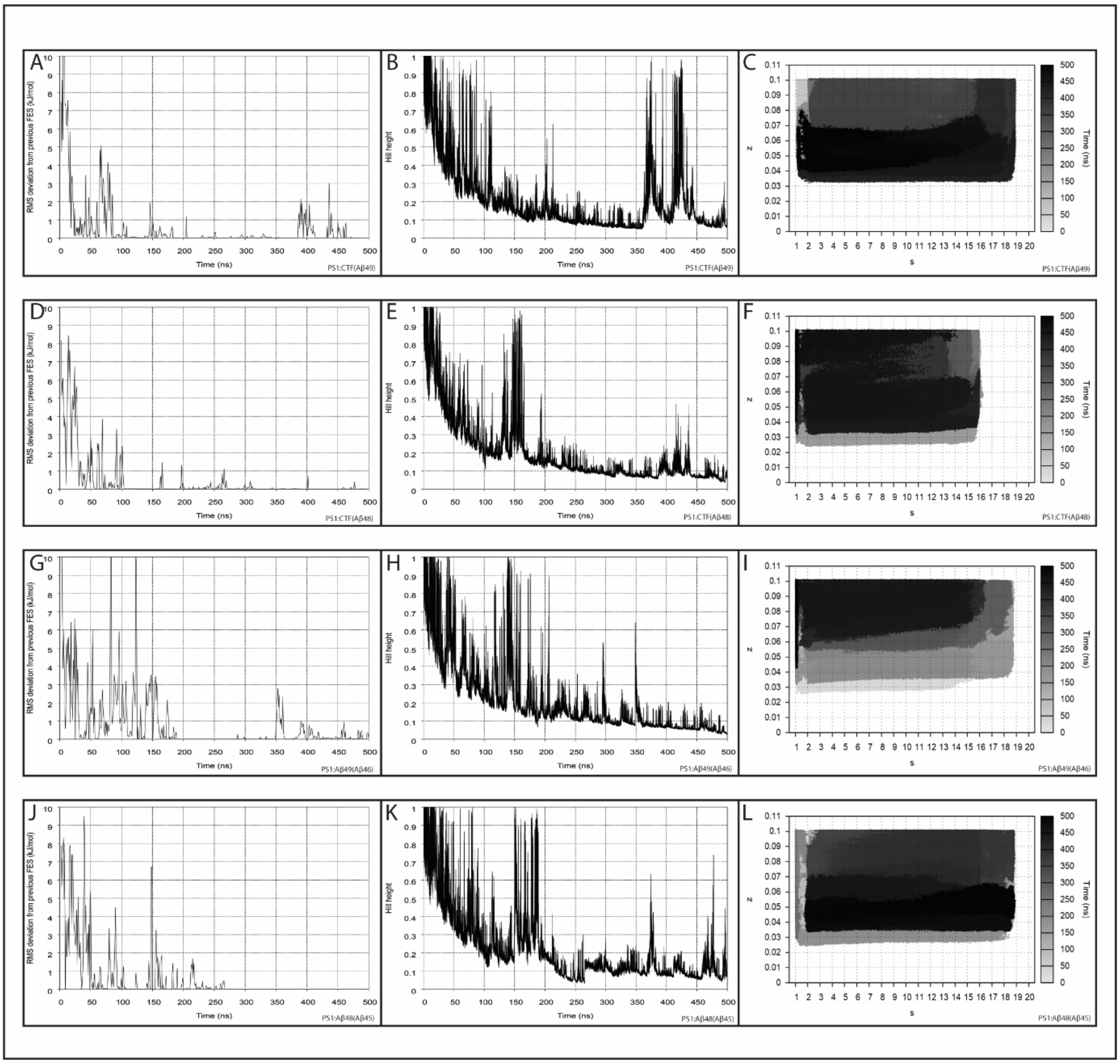
A
B
C
D
E
F
G
H
I
J
K
L
RMS deviation from previous FES (kJ/mol)
Hill height
Time (ns)
Time (ns)
s
z
PS1:CTF(Aβ49)
PS1:CTF(Aβ48)
PS1:Aβ49(Aβ46)
PS1:Aβ48(Aβ45)

**Figure S7**

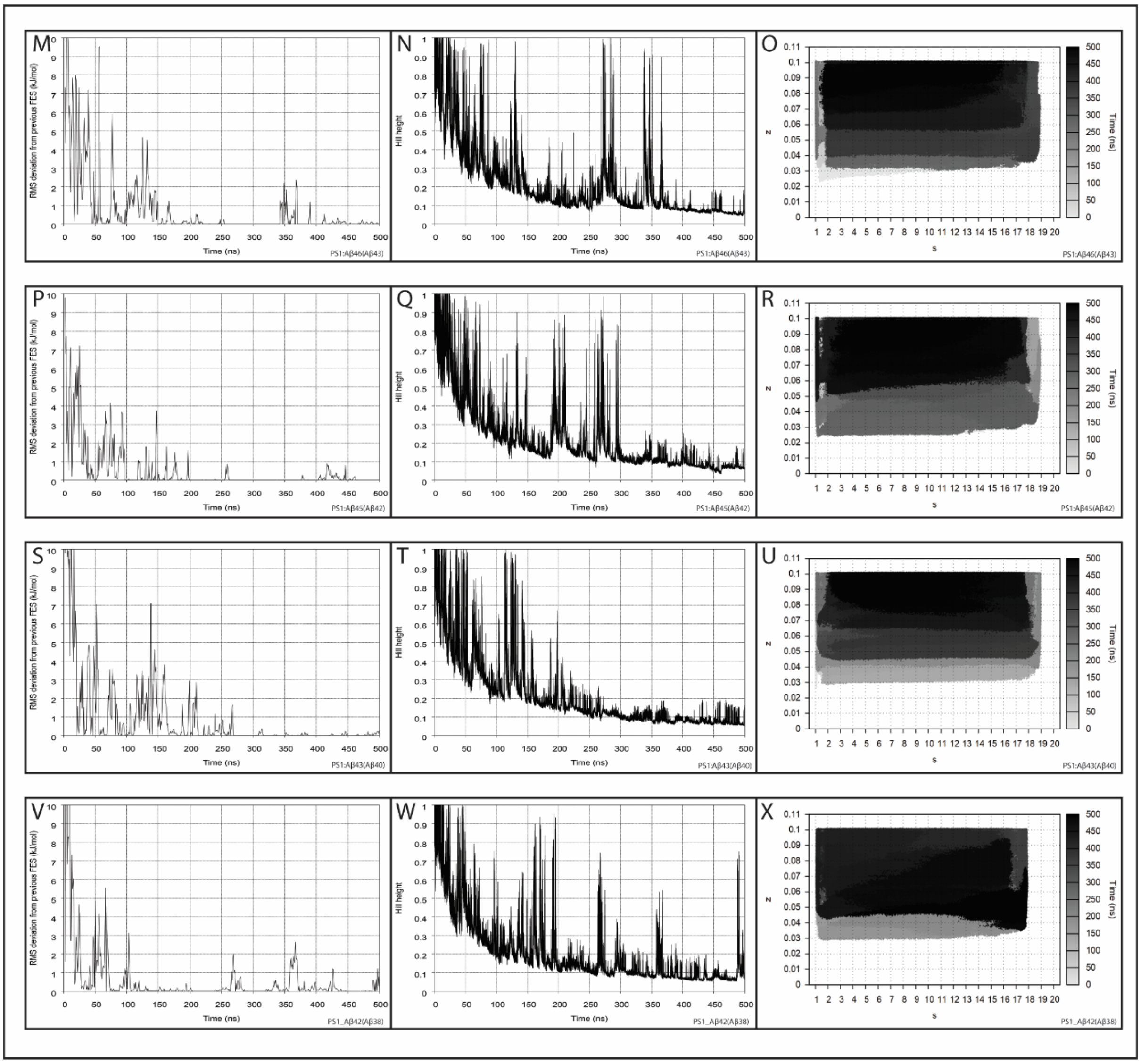


**Figure S7 continued. PS1 – APP substrate simulation convergence**. Assessed via monitoring (A, D, G, J, M, P, S, V) RMSD from previous FES at 1ns intervals, (B, E, H, K, N, Q, T, W) Gaussian hill height, and (C, F, I, L, O, R, U, X) the collective variable space over duration of simulation.

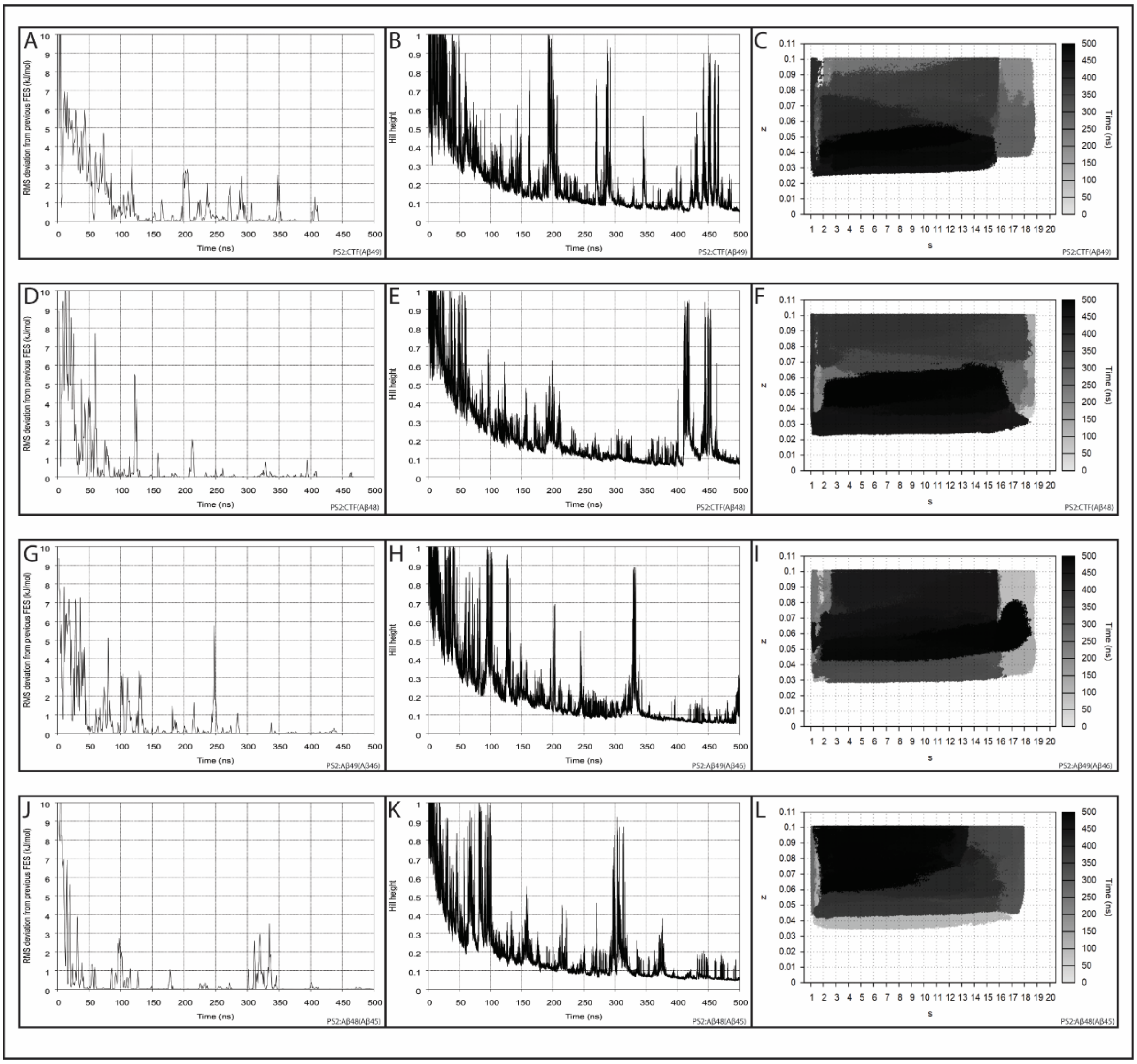
A
B
C
D
E
F
G
H
I
J
K
L
RMS deviation from previous FES (kJ/mol)
Time (ns)
Hill height
z
s
Time (ns)
0 50 100 150 200 250 300 350 400 450 500
1 2 3 4 5 6 7 8 9 10 11 12 13 14 15 16 17 18 19 20
PS2:CTF(Aβ49)
PS2:CTF(Aβ48)
PS2:Aβ49(Aβ46)
PS2:Aβ48(Aβ45)

**Figure S8**

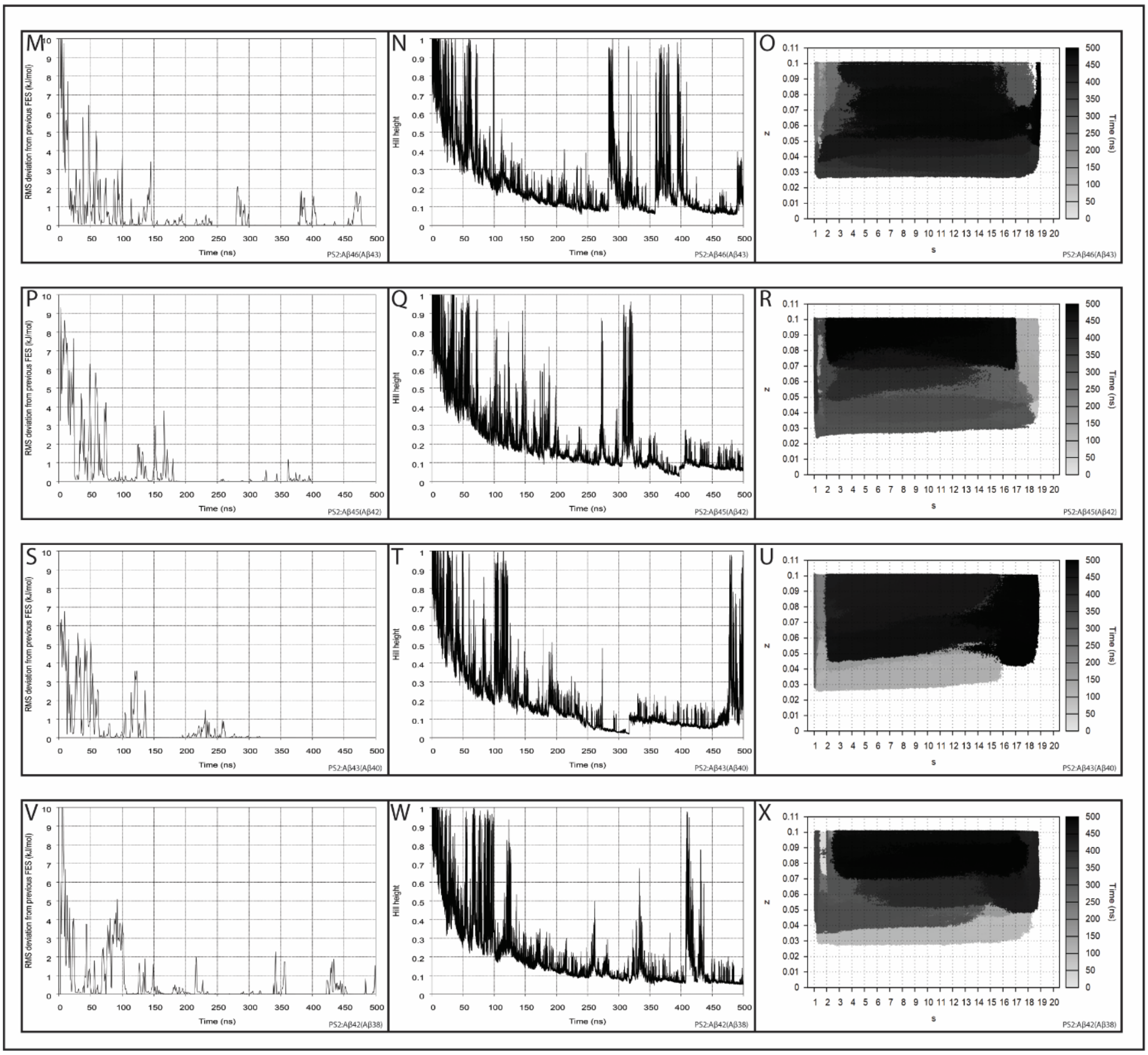

**Figure S8 continued. PS2 – APP substrate simulation convergence.** Assessed via monitoring (A, D, G, J, M, P, S, V) RMSD from previous FES at 1ns intervals, (B, E, H, K, N, Q, T, W) Gaussian hill height, and (C, F, I, L, O, R, U, X) the collective variable space over duration of simulation.

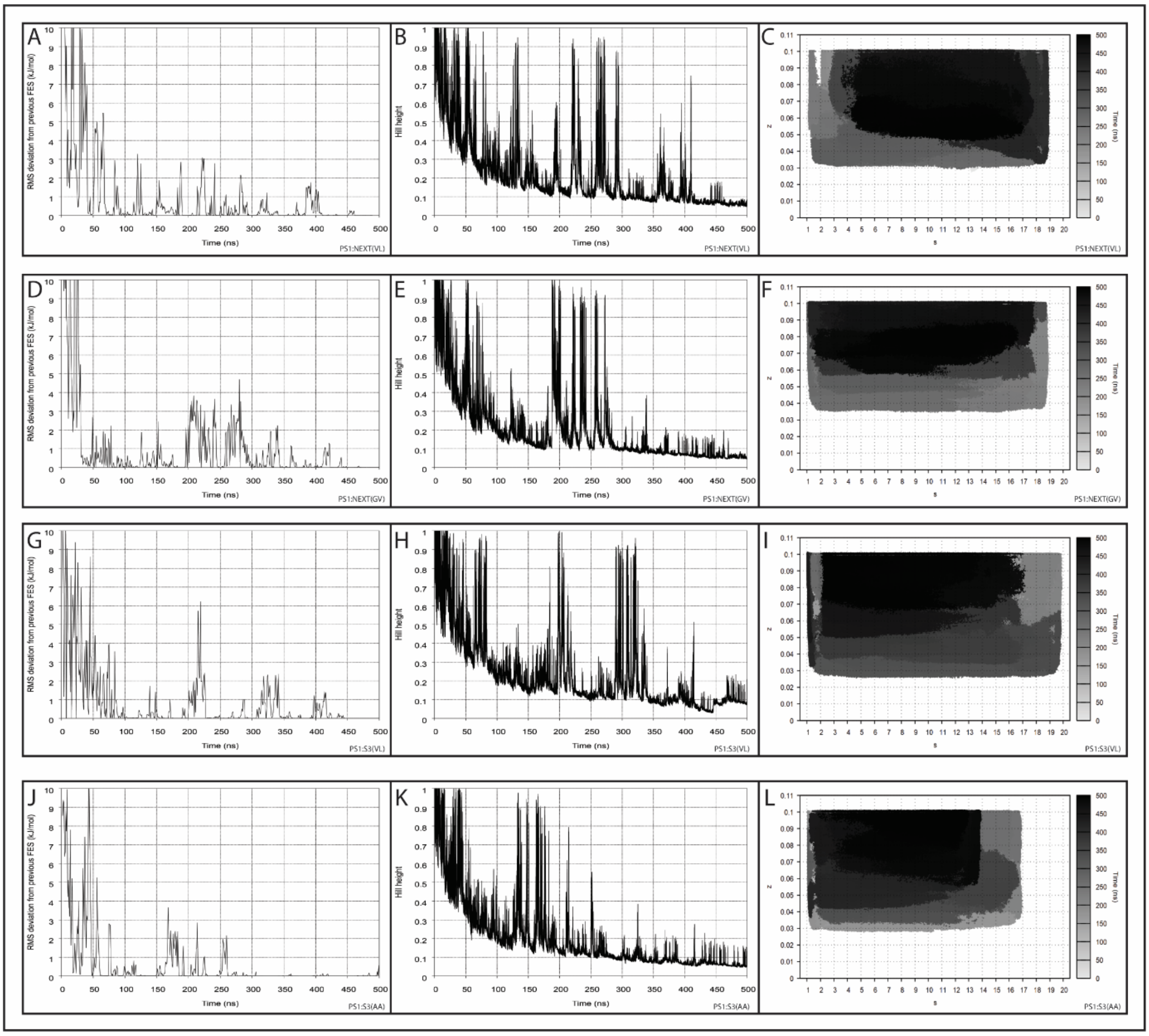


**Figure S9 PS1 – Notch substrate simulation convergence.** Assessed via monitoring (A, D, G, J) RMSD from previous FES at 1ns intervals, (B, E, H, K) Gaussian hill height, and (C, F, I, L,) the collective variable space over duration of simulation.

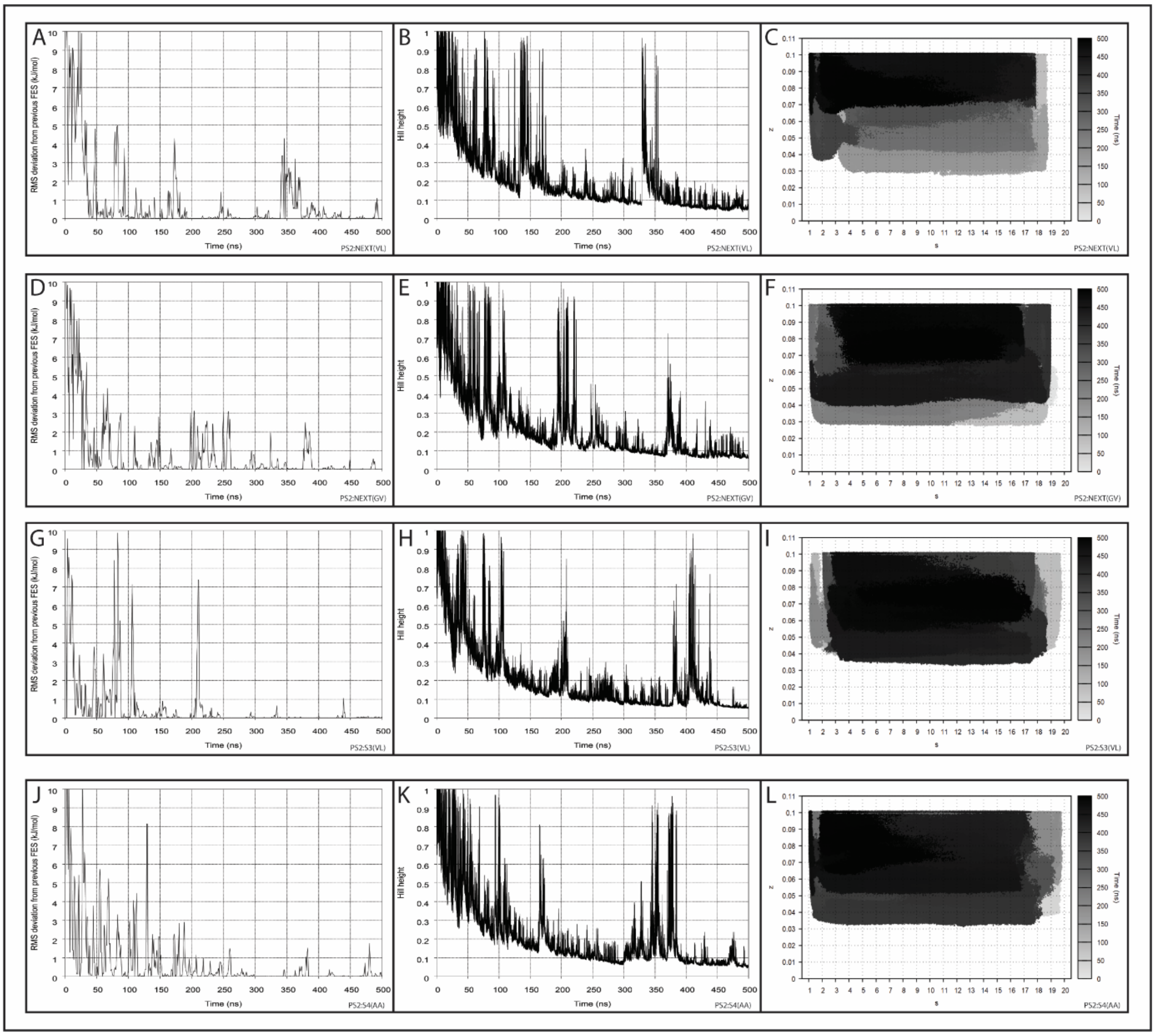


**Figure S10 PS2 – Notch substrate simulation convergence**. Assessed via monitoring (A, D, G, J) RMSD from previous FES at 1ns intervals, (B, E, H, K) Gaussian hill height, and (C, F, I, L,) the collective variable space over duration of simulation.
